# Tetramerization and RNA-guided filament assembly control Schlafen–Argonaute antiphage defense

**DOI:** 10.64898/2026.05.29.728809

**Authors:** Anton Kuzmenko, Kate Radford, Anastasiya Oguienko, Lidiya Lisitskaya, Daria Gelfenbein, Tsui-Fen Chou, Andrey Kulbachinskiy, Alexei A. Aravin

## Abstract

Two deeply conserved protein families, Argonaute (Ago) and Schlafen (SLFN), play defense roles in diverse prokaryotes and eukaryotes, including humans. Here, we identify a monophyletic group of proteins broadly distributed across bacteria and archaea that fuse a SLFN domain with an Ago core and a GHKL-family ATPase. Using SLFN-pAgo from *Runella zeae* as a model, we show that these proteins protect bacterial cells against bacteriophages by employing the Ago core as a guide-dependent sensor and the SLFN domain as a nuclease effector to induce abortive infection via tRNAs cleavage. Structural and biochemical analyses reveal that ATP binding by the GHKL domain drives tetramerization of SLFN-pAgo, reconstituting the canonical SLFN nuclease architecture found in mammalian proteins. Prior to target detection, non-canonical guide RNA binding induces the formation of long helical filaments, locking the SLFN domains in an inactive configuration. Guide-dependent recognition of complementary target DNA triggers massive structural rearrangements leading to filament disassembly and the induction of SLFN nuclease activity. Together, our findings uncover a new antiphage system that employs reversible guide- and ATP-mediated oligomerization to strictly regulate cooperation between an Ago sensor and a SLFN RNAse effector.

## INTRODUCTION

Bacteria and archaea live under relentless threat from phages and mobile genetic elements, driving the evolution of diverse defense strategies^1^. Alongside restriction–modification and CRISPR–Cas systems, prokaryotic genomes encode multiple nucleic acid–targeting defense systems whose mechanisms remain poorly understood^2–4^. Few of the multitude of prokaryotic defense proteins are conserved in eukaryotes, including Argonaute (Ago) and Schlafen (SLFN) proteins, both of which have retained central roles in genome defense across the eukaryotic tree of life^5–7^.

Argonautes use short nucleic acid guides to recognize complementary targets^8,9^. In eukaryotes, this process underlies RNA interference, miRNA-guided regulation of gene regulatory networks and piRNA-guided repression of transposable elements^10–14^. Similar to eukaryotic Agos, prokaryotic Agos (pAgos) harbor core domains including the N-terminal domain involved in guide-target interactions, MID and PAZ domains binding the 5′ and 3′ ends of the guide, respectively, and the PIWI domain mediating guide-directed target cleavage^15–18^. However, compared to their eukaryotic counterparts, pAgos exhibit striking diversity in both architecture and activity^19,20^. Only one out of three major clades of pAgos, Long-A, includes proteins that contain a catalytic tetrad in the PIWI domain and act as guide-directed nucleases^21–23^. All members of the two other clades, Long-B and short pAgos, lack the nuclease activity. Instead, such inactive pAgos were shown to act as sensors that upon detection of a complementary target activate effector domains located on the same or a separate protein^19,24^. Short pAgos rely on partner proteins containing an APAZ domain fused to various effectors, including NADases and nucleases^25–28^. Long-B pAgos also cooperate with distinct effector proteins^29^. In both groups, effector domains often mediate abortive infection, illustrating how the Argonaute platform can use different auxiliary modules to diversify immune outputs. Yet despite the prevalence of unusual domain associations, the physiological roles and molecular mechanisms of most pAgos remain poorly understood.

Similarly to Agos, SLFN proteins are broadly distributed across all domains of life and are involved in cell defense in mammals and bacteria^30–32^. First identified in mammals as regulators of thymocyte growth^33^, SLFN proteins were later shown to restrict viral infection^34^. The anti-viral activity of mammalian SLFNs relies on cleavage of tRNA or rRNA by an endonuclease site located in the SLFN domain, thus preventing virus development in infected cells^35–37^. Proteins harboring homologous SLFN domains are also widespread in bacteria and archaea, where they occur as stand-alone proteins or as fusions to diverse partners such as NACHT ATPases^30,38^. Genomic associations of SLFN in prokaryotic genomes with defense islands suggest their potential immune functions, but their contribution to prokaryotic antiviral defense has remained largely untested.

Here, we investigate the intersection of Agos and SLFNs, identify a group of SLFN-containing pAgos, define their evolutionary distribution, and explore their molecular mechanism using RzAgo from *Runella zeae* as a model. We show that pAgo core, SLFN, and a regulatory GHKL ATPase domains cooperate to control nuclease activity by switching between two functionally opposing oligomeric states — an inactive filament and an active tetramer — regulated by guide RNA loading, target DNA recognition, and ATP binding. We further demonstrate that this system provides robust protection against bacteriophages depending on its ATPase and nuclease activities and propose the mechanistic basis of this defense.

## RESULTS

### Extra-long pAgos with a conserved Schlafen domain

Phylogenetic analysis of pAgo proteins identified a novel monophyletic group that is substantially longer than other pAgos (Fig. 1a,b). Belonging to the long-B clade, this group comprises 78 unique proteins—found in both complete genomes and metagenomic sequences—that range from 1,071 to 1,187 residues in length. Like other long-B pAgos, these proteins harbor the core pAgo domains (PAZ, MID, and PIWI) and lack the catalytic tetrad within the PIWI domain. However, members of this group are distinguished by two additional domains at their N-terminus: an SLFN domain (AlbA_2, PFAM 04326) and a GHKL (Gyrase/Hsp90/Histidine Kinase/MutL) superfamily ATPase domain (PFAM 13749) (Fig. 1c). These extra-long pAgos, which we have designated SLFN-pAgos, are broadly distributed across prokaryotes. They span 16 bacterial phyla showing notable enrichment in Cyanobacteria and Bacteroidota, and include a single archaeal representative from the Asgard lineage, *Candidatus Heimdallarchaeota* (Fig. 1b). The SLFN-pAgo phylogeny is incongruent with the host species tree and contains taxonomically mixed branches, suggesting that these genes were disseminated via horizontal gene transfer (Extended Data Fig.1a).

**Figure 1.**
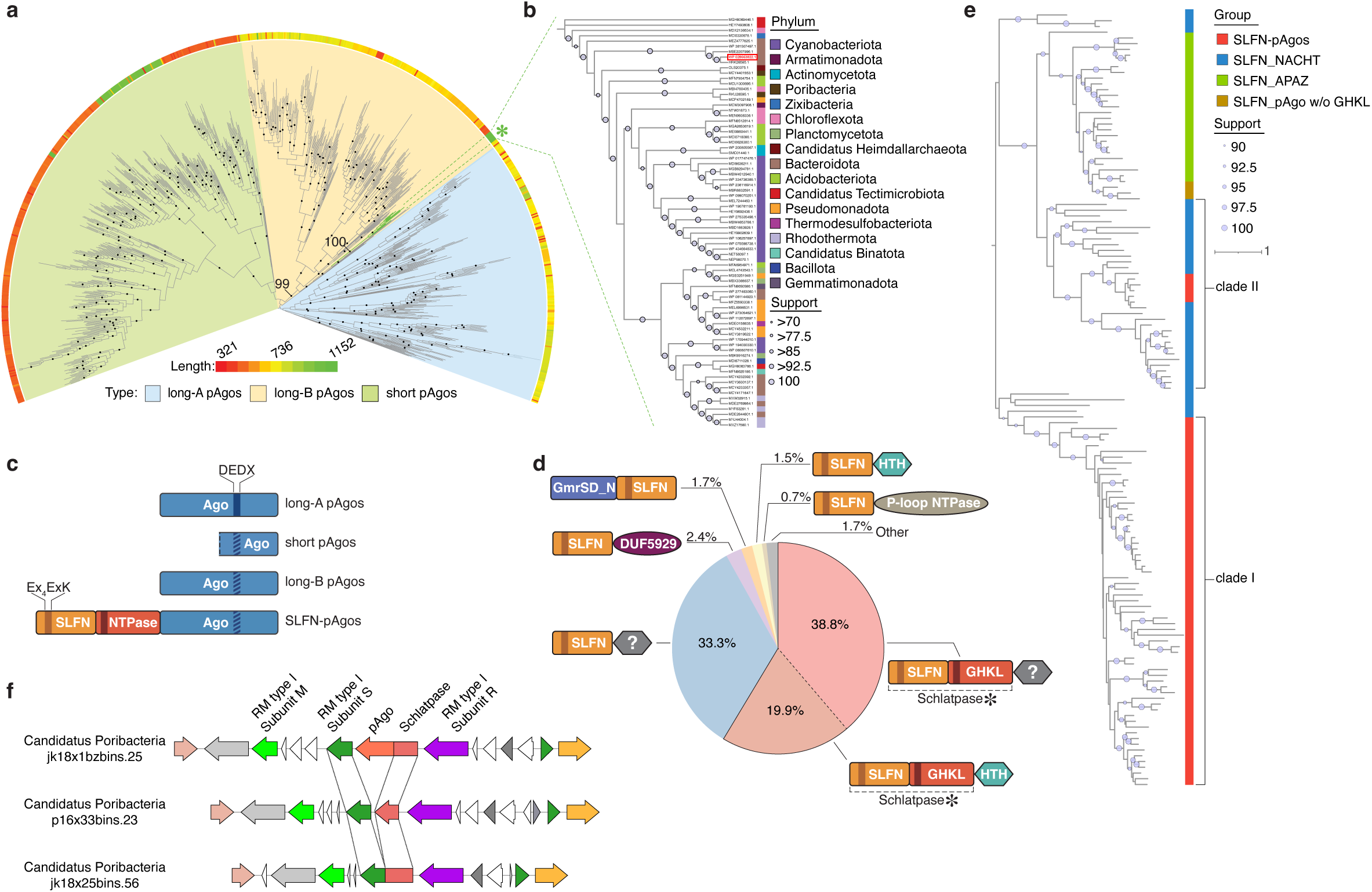
A monophyletic group of extra-long prokaryotic Argonautes is fused to a Schlatpase supradomain. **a**, Maximum-likelihood phylogeny of prokaryotic Argonautes inferred from a multiple sequence alignment of the MID and PIWI domains. The colored ring denotes protein length. Black circles mark branches supported by both ultrafast bootstrap values >95% and SH-aLRT values >80%. The asterisk indicates the SLFN-pAgo clade within long-B pAgos. **b**, Maximum-likelihood phylogeny of SLFN-pAgos inferred from a full-length multiple sequence alignment. Branches are colored by prokaryotic phylum. The SLFN-pAgo from *Runella zeae* (RzAgo; WP_028663822.1) is highlighted in red. Circle size denotes SH-aLRT support. **c**, Domain organization of SLFN-pAgos compared with the three major pAgo groups: long-A, long-B and short pAgos. The PIWI catalytic tetrad and putative active-site residues in the Schlafen_Alba2 and NTPase domains are indicated. Hatched boxes denote inactive or degenerate catalytic sites. **d**, Distribution of domain architectures among prokaryotic Schlafen-domain-containing proteins (*n* = 103,533), based on Pfam annotation. Segments outlined in black indicate proteins containing both Schlafen_Alba2 and GHKL-type NTPase domains, together forming a Schlatpase supradomain. The question mark denotes domains with no Pfam annotation. **e**, Maximum-likelihood phylogeny of Schlafen_Alba2 domains from SLFN-pAgos (red), SLFN–NACHT proteins (blue), SLFN–APAZ proteins associated with short pAgos (green) and pAgos containing Schlafen_Alba2 but lacking a GHKL NTPase domain (brown). Circle size denotes ultrafast bootstrap support. **f**, Comparative analysis of homologous operons centered on genes encoding Schlatpase-containing proteins, shown in red, from three Poribacterial species. Homologous or functionally related proteins, based on sequence annotation, are colored identically. Dotted lines trace the positions of the Schlatpase supradomain and the type I restriction–modification substrate-recognition subunit in related proteins.

Genomic analysis revealed that SLFN-pAgos are frequently encoded near defense system genes, including restriction–modification, toxin–antitoxin and BREX modules as well as phage- and transposon-related integrases (Extended Data Fig. 2). The association of SLFN-pAgos with genomic “defense islands”^39^ suggests a role in bacterial defense. However, we observed no conserved genetic linkage with specific partners, indicating that SLFN-pAgos either function autonomously or cooperate with diverse defense systems.

To investigate the origin of the distinct SLFN-pAgo domain architecture, comprising a combination of SLFN and a GHKL ATPase domains, we performed extensive sequence search in the NCBI non-redundant protein database. This analysis showed that the SLFN–GHKL ATPase combination is widespread in prokaryotes and includes multiple additional groups beyond SLFN-pAgos. Among a non-redundant set of 103,533 prokaryotic proteins containing a SLFN domain, 60,692 (58.6%) also carry the GHKL ATPase domain, suggesting their evolutionarily association (Fig. 1d). We designated this stable domain combination — a “supra-domain”^40^ — as Schlatpase (derived from *<u>Schla</u>fen–A<u>TPase</u>*). While some Schlatpase-containing proteins, including SLFN-pAgos, possess additional extended domains, the majority are smaller proteins that contain no other domains or have a short helix–turn–helix (HTH) domain at the C-terminus (Extended Data Fig. 1b).

To investigate the evolutionary relationship between SLFN domains in pAgos and other prokaryotic proteins, we reconstructed an SLFN phylogeny (Extended Data Fig. 1c, see Materials and Methods). This analysis revealed that SLFN domains of SLFN-pAgos belong to two distinct clades (Fig. 1e). Clade I domains are found almost exclusively in SLFN-pAgos, whereas Clade II domains are shared between SLFN-pAgos and members of the NACHT protein family implicated in antiviral defense^38^ suggesting that the SLFN domain was exchanged between pAgos and NACHT families. SLFN domains from Clade II are distantly related to those found in several APAZ-domain proteins associated with short pAgos^19,24^.

During a search of additional metagenomic sequences, we identified a defense operon in *Poribacteria* that encodes both a SLFN-pAgo and a restriction-modification (RM) system. Remarkably, we also found two homologous operons that lack the SLFN-pAgo. Instead, these operons contain a highly similar (>60% identity) Schlatpase module that is either fused to the substrate-recognition subunit of a type I RM system or present as a stand-alone gene (Fig. 1f). This arrangement suggests a recent exchange of the Schlatpase module between the SLFN-pAgo and RM systems. Taken together, our genomic analyses indicate that the SLFN domain is widespread across prokaryotic proteins and exists in diverse architectures, with a specific clade of this domain actively exchanged among pAgos and two other defense systems: NACHT and RM.

### RzAgo binds small RNA guides and recognizes DNA targets

To explore the structural and functional organization of SLFN-pAgos and to investigate their role in phage defense, we selected SLFN-pAgo from *Runella zeae*, RzAgo, for experimental validation. We expressed and purified RzAgo from *E. coli* (Extended Data Fig. 3a, b), evaluated its properties through a series of *in vitro* assays and determined its structure in the apo state as well as in complexes with guides and targets.

Unlike eukaryotic Argonautes, which exclusively use RNA guides to target complementary RNAs, pAgos exhibit broader guide and target specificities. To determine the guide specificity of RzAgo, we first performed *in vitro* binding assays using purified RzAgo and synthetic 18-nt single-stranded RNA or DNA with varying 5′-terminal nucleotides (Extended Data Fig. 3c). RzAgo bound RNAs with substantially higher affinity than DNAs and exhibited a strong preference for a 5′-terminal adenine (Extended Data Fig. 3c, top). Deletion of the SLFN and ATPase domains (residues 1–442) did not significantly affect the binding of short RNAs, indicating that these domains are dispensable for guide loading (Extended Data Fig. 3c, bottom). We next tested the binding of RzAgo–guide RNA complexes to complementary nucleic acids and found that these complexes preferentially bound complementary ssDNA, with negligible affinity for RNA targets (Extended Data Fig. 3d). Furthermore, deletion of the SLFN and ATPase domains of RzAgo did not affect target ssDNA binding.

To determine the guide specificity of RzAgo in bacterial cells, we purified the nucleic acids associated with it during expression in its native host *R. zeae* (see Materials and Methods). RzAgo bound to short RNA species (∼18–23 nt, median length of 21 nt) exhibiting a strong preference for 5’-terminal adenine (95.1%), consistent with our *in vitro* binding experiments (Extended Data Fig. 3e). Together these results indicate that RzAgo utilizes RNA guides to recognize complementary DNA targets, and that neither guide binding nor target recognition depends on the Schlatpase supradomain.

### Loading of guide RNA drives formation of RzAgo filaments

To gain insight into the structural organization of RzAgo, we determined the cryo-EM structure in the absence of guide or target at an overall resolution of 3.48 Å (Fig. 2a, Extended Data Fig. 10). No clear density corresponding to the Schlatpase supradomain was observed, suggesting its high conformational heterogeneity and/or partial instability (Extended Data Fig. 4c). Indeed, AlphaFold3 predicts two flexible linkers connecting the SLFN and ATPase domains and the ATPase and pAgo core domains (Extended Data Fig. 4a, b). The resolved portion of RzAgo adopts the typical bilobal architecture of Argonaute proteins, with a notable absence of the N-terminal domain characteristic to most long pAgos. Consequently, the nucleic acid–binding cleft is formed between two asymmetric lobes, a larger lobe comprising the MID and PIWI domains and a smaller lobe composed of the L1 and PAZ domains. In the absence of guide RNA, these domains appear conformationally heterogeneous, as reflected by a gradual loss of resolution toward solvent-exposed regions, suggesting intrinsic flexibility of the pAgo core prior to guide loading (Extended Data Fig. 10).

**Figure 2.**
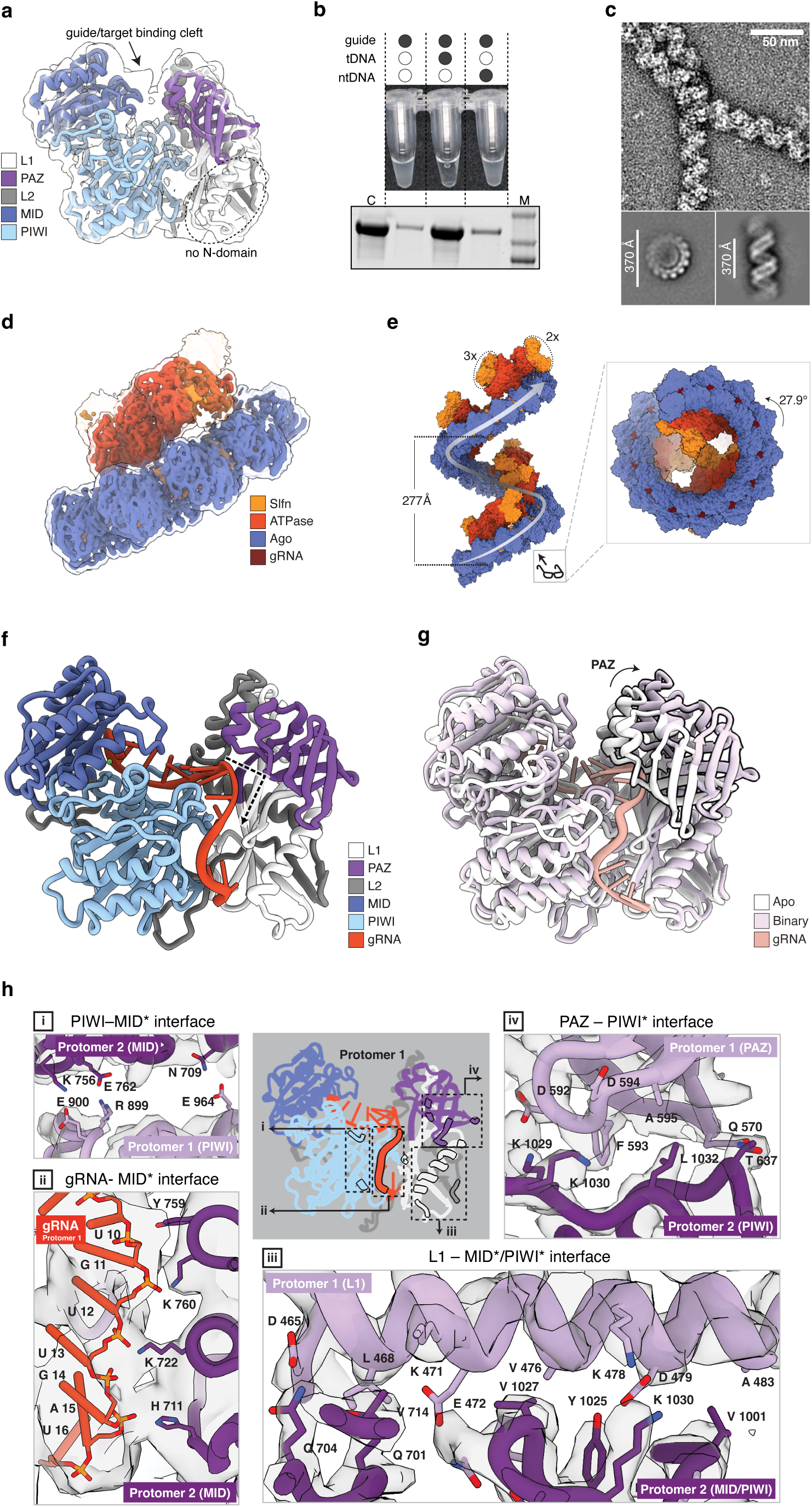
Guide RNA binding induces reversible oligomerization of RzAgo. **a**, Cryo-EM model of apo RzAgo fitted into a low-contour density map, shown as a transparent surface. pAgo core domains are color-coded. The dashed oval indicates the position of the absent canonical N-terminal domain. The guide and target-binding cleft is indicated by an arrow. **b**, Top, guide RNA induces visible precipitation of RzAgo, which is reversed by addition of a fully complementary DNA target. Bottom, SDS–PAGE analysis of soluble RzAgo remaining in the supernatant after incubation with guide RNA alone, with a fully complementary target DNA (tDNA), or with a non-complementary DNA control (ntDNA). **c**, Top, representative negative-stain EM image of an RzAgo filament formed in the presence of guide RNA. Bottom, representative two-dimensional class averages showing top and side views of the helical filament. **d**, Sharpened cryo-EM model of the RzAgo filament comprising five consecutive protomers, obtained after mild crosslinking. Protein domains and guide RNA are color-coded; low-contour transparent density highlights the positions of Schlafen domains not visible in the sharpened map. **e**, Left, composite space-filling model of the RzAgo helical filament built from the cryo-EM reconstruction in **d**. Right, axial view from below. Helical parameters are indicated. **f**, Structure of an individual RzAgo monomer, highlighting a pronounced kink in the bound guide RNA, indicated by a dashed arrow. **g**, Superposition of the pAgo core domains in apo and binary RzAgo. The arrow indicates guide-RNA-induced movement of the PAZ domain, outlined in black. **h**, Contacts between the pAgo core domains of two consecutive protomers, designated 1 and *. The central panel shows protomer 1, with structural elements involved in the intersubunit interface outlined in black. Arrows indicate magnified views of protein–protein contacts (**I**, **III** and **IV**) and RNA–protein contacts (**II**) that stabilize intersubunit interactions within the filament.

We next sought to determine whether guide RNA and target DNA binding could induce structural changes in RzAgo. Upon incubating guide RNA with RzAgo, we observed that the solution became opaque within minutes, indicating potential multimerization of the protein-RNA complexes (Fig. 2b, top). This corresponded to a large decrease in the amount of soluble RzAgo (Fig. 2b, bottom). Subsequent addition of single-stranded target DNA complementary to the guide RNA — but not non-target DNA — restored solution clarity. This suggests that target DNA binding induces the disassembly of multimeric structures formed by RzAgo binary complexes.

To visualize RzAgo-RNA multimers, we employed negative-stain electron microscopy (ns-EM). While apo-RzAgo appeared as monodisperse particles, RzAgo–RNA complexes assembled into regular right-handed helical filaments up to several hundred nanometers in length. This indicates that interaction with guide RNA triggers the orderly multimerization of RzAgo rather than non-specific aggregation (Fig. 2c, Extended Data Fig. 4d). The subsequent addition of complementary target DNA to pre-assembled RzAgo-RNA complexes led to disassembly of these filaments back into monodisperse particles.

To understand the molecular mechanism of filament formation, we analyzed the structure of RzAgo with an 18-nt guide RNA using cryo-EM. The complexes were obtained with or without mild glutaraldehyde crosslinking to stabilize their conformation, and corresponding structures were solved to a nominal resolution of 4.4 Å and 4.2 Å, respectively (Fig. 2d, Extended Data Fig. 4e). Using crosslinking, we obtained a structure comprising five consecutive RzAgo protomers, which we used to generate a composite filament model assembled from repeating units. In agreement with ns-EM, the composite model revealed that RzAgo-RNA complexes form right-handed helical filaments with ∼12.5 subunits per turn, with a rise of 21.5 Å, twist of 27.9° per protomer, and pitch of 277 Å (Fig. 2e). The backbone of the filament is formed by pAgo core domains (PAZ, MID and PIWI), which are assembled in a head-to-tail arrangement, while the SLFN and ATPase domains are visible only with glutaraldehyde crosslinking.

Similar to other Ago proteins, the 5′ end of the guide RNA is anchored within a conserved pocket in the MID domain of RzAgo. Here, specific contacts between the 5’-adenine and residues Y828 and Y1086 explain the strong preference for adenosine at the first guide position (Extended Data Fig. 5a–c). In contrast, the binding of the 3′ portion of the guide RNA in RzAgo differs markedly from that observed in known pAgos. Instead of a canonical N domain, which directs the guide toward the PAZ domain in other pAgos, RzAgo and other SLFN-pAgos possess a patch of acidic residues within the L1 and PAZ domains (Extended Data Fig. 5d, top). Driven by electrostatic repulsion from this patch, the 3′ segment of the guide is rerouted into a channel formed between the L1 and PIWI domains of one protomer and the MID and L2 domains of the adjacent protomer (Extended Data Fig. 5e). To adopt this non-canonical trajectory, the RNA makes a ∼90° turn around nucleotides 9 and 10, which is stabilized by a series of hydrogen bonds mediated by the PIWI and L1 domains (Fig. 2f).

Comparison of the apo and binary states of RzAgo reveals that guide binding induces a ∼10 Å separation between the L1/PAZ and MID/PIWI lobes. This separation repositions the PAZ domain and widens the inter-lobe cleft to accommodate the guide RNA (Fig. 2g; Extended Data Fig. 6). The conformational rearrangements triggered by RNA binding enable extensive interprotomer contacts, mediated by a combination of protein–protein and RNA-mediated interactions. Specifically, movements of several loops in the MID and PIWI domains bring together the L1 helix, PAZ domain, PIWI residues, and 3′-end of the guide RNA of one protomer with the L2 domain, hinge-forming PIWI loops, and a cluster of basic residues on the outer surface of the neighboring protomer’s MID domain. Consequently, the PIWI domain contacts the MID domain of the adjacent protomer (Fig. 2h-i), the long helix in the L1 linker engages regions of the adjacent MID and PIWI domains (Fig. 2h-iii), and the PAZ domain interacts with the neighboring PIWI domain (Fig. 2h-iv) as well as with portions of its L2 linker and MID domain. Furthermore, the non-canonical binding of the guide’s 3’ end enables the RNA to directly contact surface-exposed residues on the MID domain of the neighboring protomer (Fig. 2h-ii, Extended Data Fig. 5d, bottom). Thus, the guide RNA plays an essential role in stabilizing RzAgo filaments, both supporting an interface-forming conformation and acting as a molecular ‘glue’ between adjacent protomers. Overall, the binary structure elegantly explains why filament formation is induced by guide loading.

### DNA target recognition reroutes the guide RNA and destabilizes the RzAgo filament

To gain mechanistic insight into how target DNA binding induces RzAgo filament disassembly, we prepared a ternary complex comprising RzAgo, an 18-nt 5′-phosphorylated guide RNA, and a complementary 24-nt DNA. Single-particle cryo-EM analysis reconstructed the ternary complex structure with a nominal resolution of 3.3 Å (Fig. 3a, Extended Data Fig. 10). In the resulting model, the RNA–DNA heteroduplex occupies the cleft between the L1/PAZ and MID/PIWI lobes and is positioned through an extensive network of hydrogen bonds formed primarily with the sugar–phosphate backbone (Extended Data Fig. 5h). It is fully base-paired beginning at the second position of the guide (U2) and adopts a non-canonical geometry distinct from both A- and B-form helices—a feature previously observed in the RsAgo ternary complex^41^ (PDB: 6D8P, Extended Data Fig. 5i). The first nucleotide of the guide (A1) and its corresponding target base (dT21) are splayed apart and remain unpaired. Despite lacking an N domain, RzAgo positions the distal end of the heteroduplex in essentially the same location as RsAgo, indicating that the N domain is not required for heteroduplex positioning.

**Figure 3.**
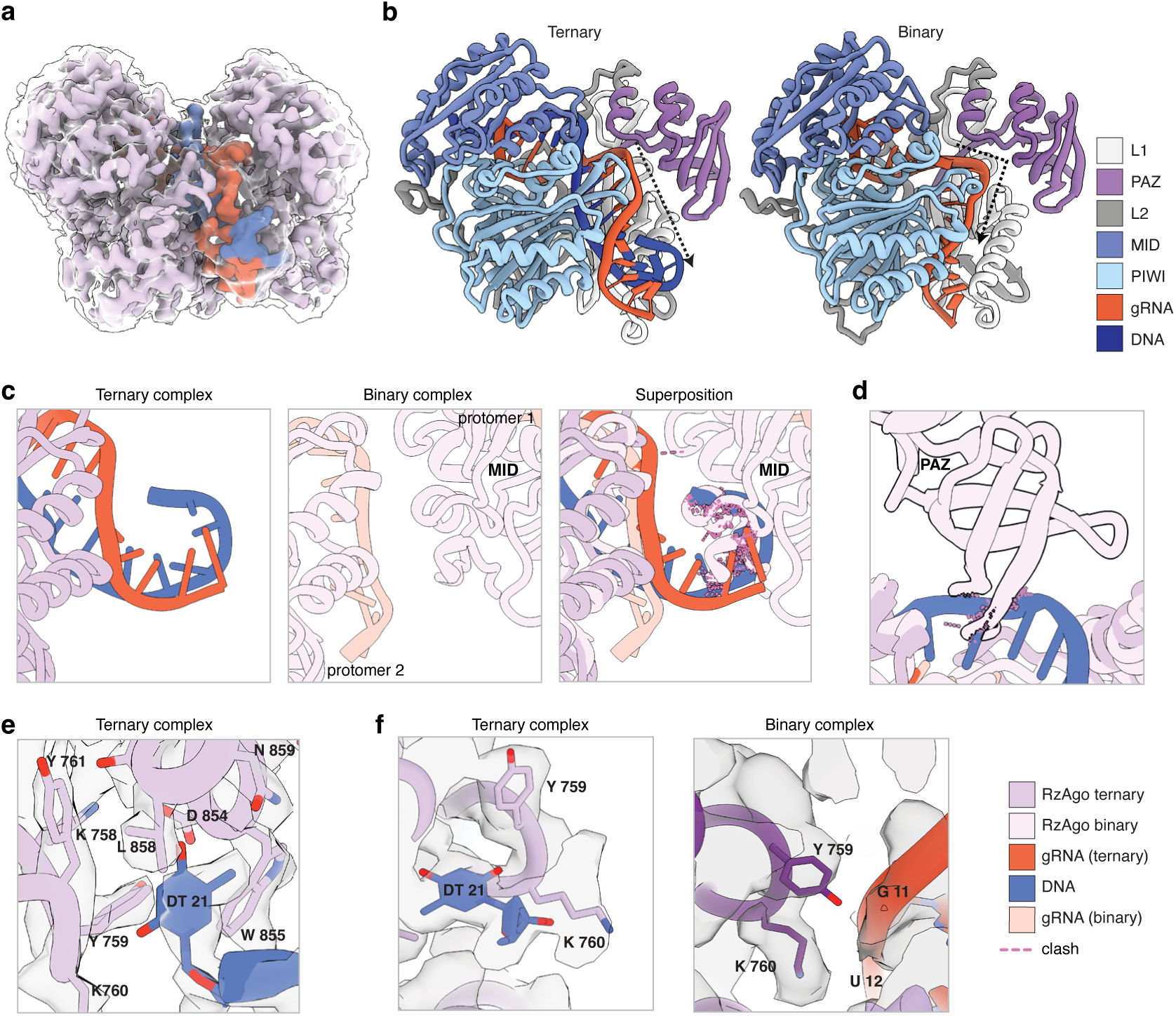
Structural basis for target-DNA-induced disassembly of RzAgo filaments. **a**, Sharpened cryo-EM map of RzAgo bound to an 18-nt guide RNA and a fully complementary 24-nt target DNA, overlaid with a low-contour density map shown as a transparent surface. Guide RNA and target DNA are shown in red and blue, respectively. **b**, Comparison of the guide RNA trajectory in binary and ternary RzAgo complexes. The dashed line highlights the change in trajectory associated with formation of the RNA–DNA heteroduplex. **c**, Superposition of ternary and binary RzAgo complexes, revealing a steric clash between the distal end of the RNA–DNA heteroduplex and a neighboring RzAgo protomer in the filament. **d**, Steric clash between the PAZ domain of the neighboring protomer and the target DNA strand upon heteroduplex formation. **e**, Surface pocket in the MID domain that engages nucleotide dT21 of the target DNA. **f**, Structural comparison of the dT21-binding pocket in the binary and ternary states, showing conformational rearrangement of pocket residues. Protein domains, guide RNA, target DNA and steric clashes are color-coded as in **c–f**.

In the ternary complex, base-pairing between the guide RNA and target DNA alters the RNA trajectory, preventing its interactions with the preceding protomer (Fig. 3b). Instead of bending, the RNA–DNA duplex spans the cleft along its longest path and sterically blocks the filament-forming interface, clashing with interacting residues of the MID domain of the neighboring protomer (Fig. 3c). Furthermore, the target DNA strand occupies the same space as the neighboring PAZ domain in the binary complex multimer, producing a steric clash incompatible with the filament arrangement (Fig. 3d). The target DNA thymine complementary to the 5’-adenine of guide RNA (dT21) is flipped away and locked in a pocket on the surface of the MID domain (Fig. 3e). Notably, several residues that mediated interprotomer contacts in the filament now contribute to this pocket (Fig. 3f). Together, these data explain why target DNA binding is incompatible with filament formation.

Target binding drives the coordinated rearrangement of a protein loop that connects β-strands 1 and 2 in the PIWI domain (891–903) (Extended Data Fig. 5j). In the binary complex, this loop interacts with the guide around positions 10–12, but shows weak and fragmented density indicating its substantial mobility. In the ternary complex, guide–target pairing is accompanied by a marked conformational transition, during which this loop refolds into a shortened β-hairpin that interrogates the duplex at positions U10–U12 across the minor groove. In this conformation, the PIWI loop engages both guide and target strands through backbone- and base-level contacts, suggesting that it acts as a structural element that senses duplex formation.

### ATP binding by the GHKL domain drives RzAgo tetramerization

The histidine kinase–like ATP-binding domain (PFAM 13749) in RzAgo belongs to the broader GHKL superfamily found in diverse proteins, many of which bind and hydrolyze ATP. These proteins generally function as dimers or tetramers, and the binding of ATP acts as a molecular switch, promoting the dimerization of the N-terminal ATPase domains to form a stable, closed conformation that allows for ATP hydrolysis^42,43^.

To test whether RzAgo binds nucleotides, we employed the differential radial capillary action of ligand assay (DRaCALA). We found that RzAgo binds ATP with submillimolar affinity, and that deletion of the Schlatpase supradomain abolished this interaction (Fig. 4a). Competition assays with excess unlabeled GTP, CTP, or UTP demonstrated a high specificity for ATP (Fig. 4b). Finally, thin-layer chromatography (TLC) showed that RzAgo hydrolyzes ATP to ADP and inorganic phosphate (Fig. 4c).

**Figure 4.**
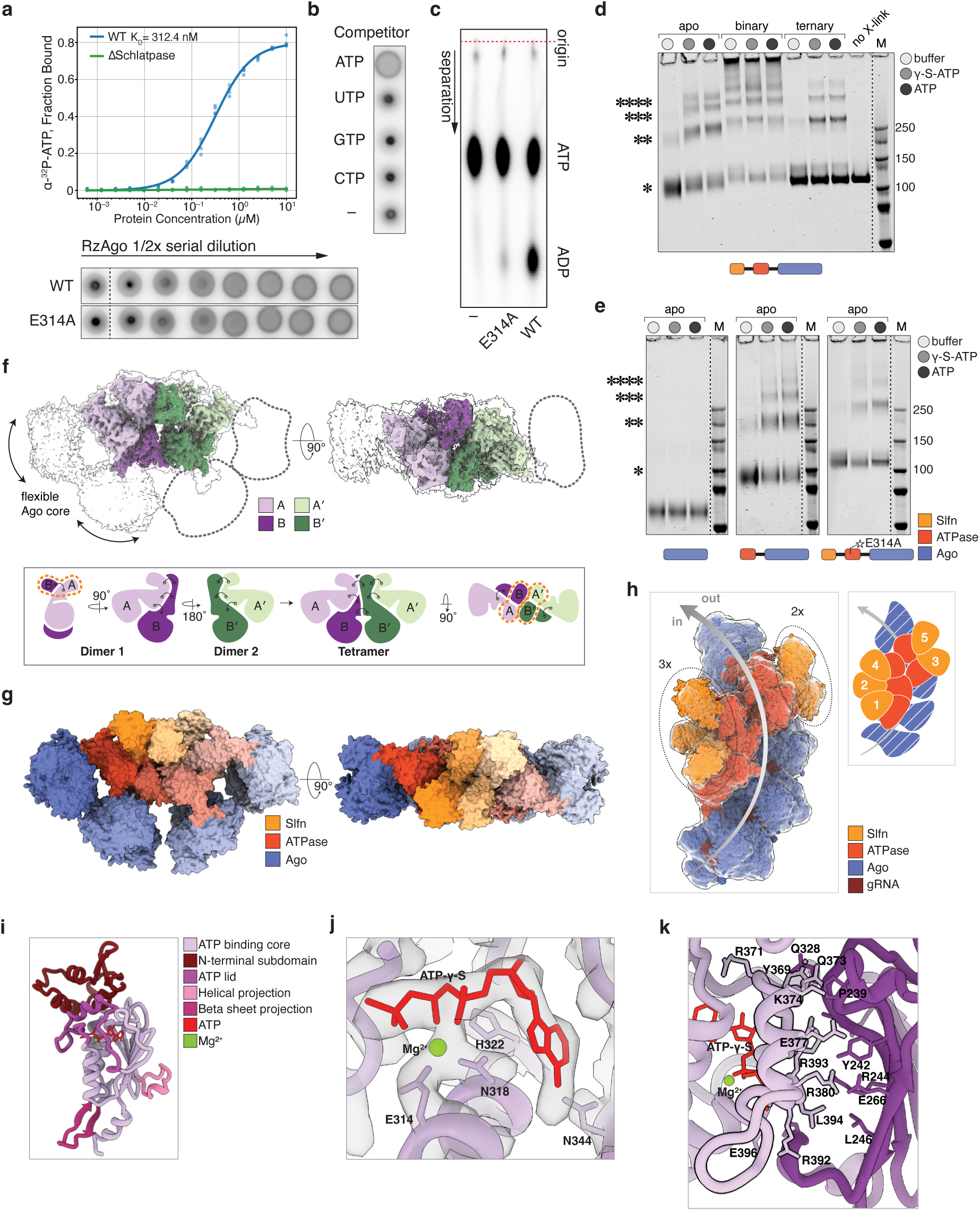
ATP binding and hydrolysis promote RzAgo tetramerization. **a**, Top, ATP-binding isotherms for wild-type RzAgo and a Schlatpase-deletion mutant. Bottom, representative DRaCALA image comparing ATP binding by the E314A mutant and wild-type RzAgo. **b**, Competition DRaCALA assay with unlabeled nucleotides, demonstrating selective ATP binding. The minus sign denotes a control without unlabeled NTPs. **c**, Thin-layer chromatography analysis of ATP hydrolysis by wild-type RzAgo and the ATPase-deficient E314A mutant. The origin marks the starting position. The minus sign denotes a control without RzAgo. **d**, Crosslinking SDS–PAGE analysis of oligomerization of wild-type RzAgo in apo, binary and ternary states. Positions of monomers, dimers, trimers and tetramers are indicated. no X-link, non-crosslinked control; M, molecular weight marker; γ-S-ATP, non-hydrolysable ATP analogue. **e**, Crosslinking SDS–PAGE analysis of RzAgo mutants, schematized below, in the apo state. Labels are as in **d**. **f**, Top, cryo-EM map of the RzAgo tetramer in side and top views, with individual protomers (A, A′, B and B′) color-coded. Low-contour transparent density highlights the positions of two unresolved pAgo core domains, which are also indicated by dashed outlines. Bottom, schematic showing hierarchical tetramer assembly from two identical dimers arranged with pseudo-C2 symmetry. Schlafen-domain dimers are indicated by dashed lines. **g**, Space-filling cryo-EM model of the RzAgo tetramer, colored by domain, shown in side and top views as in **f**. The positions of pAgo core domains were modelled from an AF3 prediction. **h**, Left, space-filling cryo-EM model of the RzAgo filament within a low-contour transparent density, showing Schlafen domains arranged into two spatially separated clusters containing two and three domains. Right, schematic showing the domain positions in the filament. Numbers indicate protomer positions within the helix. The arrow indicates the path of the helix. **i**, Structure of the RzAgo GHKL ATPase domain, with major structural elements color-coded. Bound ATP and magnesium ion are shown in red and green, respectively. **j**, Close-up view of the ATP-binding pocket in the GHKL domain, highlighting residues involved in ATP coordination and Mg²⁺ binding. **k**, Intersubunit contacts between two RzAgo monomers within the ATP-lid-mediated dimer interface. Contacting residues are highlighted, and the ATP lid is outlined in black.

To determine how ATP modulate RzAgo oligomerization, we analyzed glutaraldehyde-crosslinked RzAgo complexes using denaturing gel electrophoresis (Fig. 4d). Guide-free RzAgo lacked oligomers without ATP, whereas adding ATP or its non-hydrolyzable analog (γ-S-ATP) induced dimer, trimer, and tetramer formation. Upon adding guide RNA, large RzAgo complexes were retained in the gel wells irrespective of ATP, demonstrating that binary filament assembly is ATP-independent. While complementary target DNA dissociated these large oligomers, the dimers and tetramers remained intact. To test whether guide RNA binding is required for tetramerization, we mutated two residues in the MID domain guide-binding pocket to create RzAgo-YK. This mutant fails to bind guide RNA *in vitro* (Extended Data Fig. 5f) yet still forms tetramers upon ATP addition, confirming that guide binding is dispensable for tetramerization (Extended Data Fig. 5g).

To gain structural insight into the role of ATP binding in RzAgo oligomerization, we determined the cryo-EM structure of the RzAgo tetramer in the presence of γ-S-ATP at a 3.3 Å resolution (Fig. 4f, top; Fig. 4g; Extended Data Fig. 10). In the resulting structure, interactions between subunits are mediated by the SLFN and ATP-binding domains, while the core pAgo domains do not participate in oligomerization and are only partially resolved (Fig. 4f). In agreement with our structural analysis, deletion of the entire Schlatpase supradomain (residues 1–442) abolished tetramer formation, whereas deletion of the SLFN domain alone (residues 1–142) had only a minor effect (Fig. 4e). Thus, the ATPase domain plays the dominant role in tetramerization. Consequently, the architecture of RzAgo tetramers is drastically different from that of the filaments, in which intersubunit contacts are mediated primarily by the pAgo cores (Fig. 4h).

The RzAgo tetramer is hierarchically organized as two dimers that further associate into a higher-order assembly with C2 symmetry (Fig. 4f, bottom). Within each dimer, the two protomers adopt distinct conformations (designated A and B), which differ in the relative orientation of the SLFN and ATP-binding domains (Extended Data Fig. 7a). The dimer interface has a buried surface area of ∼1,910 Å² and is formed primarily by the ATP-binding domains (∼67% of the interface), with additional contributions from an α-helix at the base of the SLFN domain (residues 151–161, Extended Data Fig. 7b). Two dimers then associate through a larger interdimer interface (∼2,600 Å²), which is dominated by polar interactions contributed by both the SLFN (∼60%) and ATP-binding domains (Extended Data Fig. 7c), resulting in a C2-symmetric compact tetrameric architecture in which each protomer contacts two neighboring subunits via distinct domain interfaces (Extended Data Fig. 7d). The C2 symmetry of the tetramer arranges pAgo cores in an anti-parallel configuration, which is sterically incompatible with the head-to-tail packing required for filament assembly.

The ATP-binding domain of RzAgo adopts a fold closely resembling the Bergerat-type ATP-binding domain of human pyruvate dehydrogenase kinase 4: an α–β sandwich containing a seven-stranded antiparallel β-sheet and three opposing α-helices, with a loop–helix–loop ATP lid traversing the helical face (Fig. 4i, Extended Data Fig. 8a-c). The γ-phosphate and bound magnesium ion are coordinated by residues E314 and N318 (Fig. 4j). Consequently, mutation of the conserved E314 residue abolished ATP hydrolysis while preserving ATP binding (Fig. 4a, c). This mutant still formed ATP-dependent oligomers, albeit with reduced efficiency (Fig. 4e). In the ATP-bound RzAgo tetramer, many oligomerization interfaces map to conserved elements of the GHKL domain, including the ATP lid, providing a structural basis for our biochemical observation that ATP binding drives RzAgo complex formation (Fig. 4k, Extended Data Fig. 7e, f).

### RzAgo dimerization structurally mimics mammalian SLFNs to enable tRNA cleavage

The SLFN domain of RzAgo adopts a two-layer α+β topology similar to the C-lobe of mammalian SLFN domains, which harbor the nuclease active site (Fig. 5a, Extended Data Fig. 8d). Consistent with their mammalian counterparts, SLFN-pAgos feature a conserved E(x)_4_(D/E)xK nuclease motif, a SAFAN sequence (Schlafen box) and a conserved stretch of glycine and hydrophobic residues that comprise the domain core (Fig. 5a, b). Structural analysis revealed that the RzAgo catalytic pocket closely resembles that of human SLFN11, featuring a triad of acidic residues coordinating a metal ion – consistent with a divalent cation-dependent nuclease mechanism (Fig. 5c).

**Figure 5.**
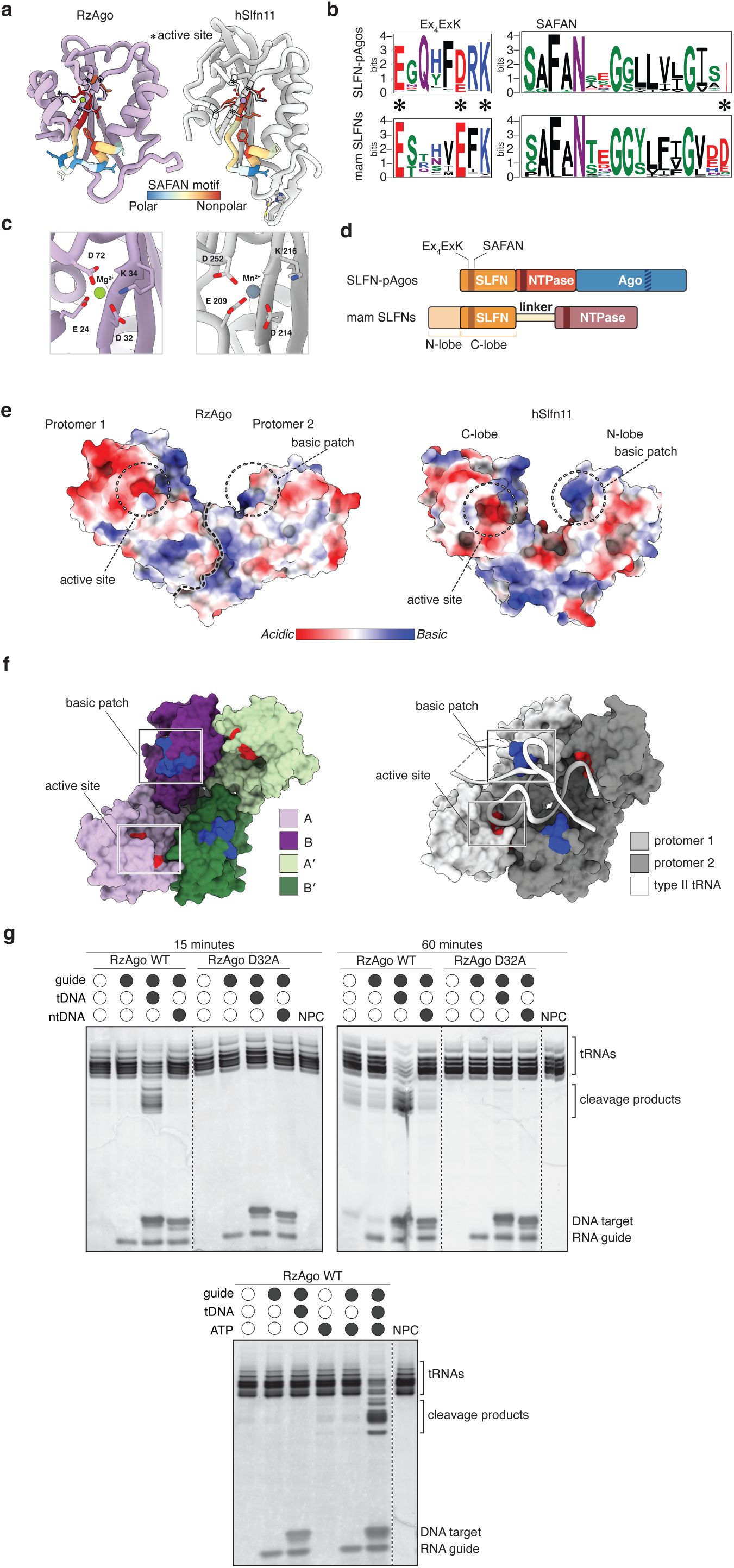
The RzAgo Schlafen domain mimics mammalian Schlafen proteins and cleaves tRNAs. **a**, Structural comparison of the Schlafen domain from RzAgo and human SLFN11 (hSLFN11). Active-site residues are marked by asterisks and outlined in bold. The highly conserved SAFAN motif and ten downstream amino acid residues are colored according to hydrophobicity. **b**, Sequence logos showing conservation of nuclease active-site residues and the SAFAN motif in SLFN-pAgos and mammalian Schlafen proteins. Asterisks mark active-site residues. **c**, Close-up views of the SLFN nuclease active sites of RzAgo and hSLFN11, showing conserved residues involved in divalent-metal-ion coordination. **d**, Domain organization of SLFN-pAgos and type III mammalian Schlafen proteins. **e**, Space-filling models of the SLFN dimer in the RzAgo tetramer and of a single SLFN domain from hSLFN11, colored by electrostatic potential. The dashed line marks the boundary between SLFN domains in RzAgo. The nuclease active site and a conserved basic surface patch are indicated. **f**, Space-filling models of SLFN domains in the RzAgo tetramer, left, and hSLFN11 dimer, right, bound to a type II tRNA substrate. Domains are color-coded by protomer. The GHKL ATPase and pAgo core domains of RzAgo, and the linker and P-loop NTPase domain of hSLFN11, are omitted in **e** and **f** for clarity. **g**, tRNA cleavage assay in the presence of ATP and wild-type RzAgo or the nuclease-deficient D32A mutant in apo, binary or ternary states. tDNA, fully complementary DNA target; ntDNA, non-complementary DNA control; NPC, no-protein control. Positions of tRNAs, cleavage products, RNA guides and DNA targets are indicated.

In mammalian SLFN domains, the nuclease C-lobe and the N-lobe together form a characteristic horseshoe-shaped architecture, with the N-lobe providing a basic patch for RNA substrate binding (Extended Data Fig. 8e)^44,45^. In contrast, SLFN-pAgos lack the N-lobe (Fig. 5d). Surprisingly, our cryo-EM model of the RzAgo tetramer revealed that two SLFN domains within a dimer assemble into a horseshoe-shaped configuration that recapitulates the bilobal organization of mammalian Schlafens (Fig. 5e). In the dimer, the catalytic pocket of one protomer faces a basic surface patch (K46, K47) on the opposing protomer, mimicking the relative positioning of functional surfaces within a single mammalian SLFN domain. Thus, dimerization of RzAgo effectively reconstructs a two-lobed SLFN architecture, with one protomer contributing the catalytic lobe and the other providing a basic surface. In human SLFN11, dimerization creates a substrate-binding cleft that accommodates type II tRNA. The tetrameric organization of RzAgo positions two dimers in a remarkably similar overall configuration, albeit with a wider cleft (Fig. 5f).

We conducted *in vitro* assays using total *E. coli* tRNA to evaluate RzAgo nuclease activity (Fig. 5g). While both apo- and guide-bound RzAgo exhibited minimal basal activity, the addition of complementary target DNA to the binary complex in the presence of ATP triggered robust tRNA cleavage. A single-residue mutation (D32) within the putative SLFN active site completely abolished this cleavage, confirming the SLFN domain as the catalytic module. These results demonstrate that guide-dependent DNA target recognition licenses the RzAgo SLFN domain to cleave tRNA.

### RzAgo protects cells from phage infection

To explore the ability of RzAgo to act in defense against phages we used a related Bacteroidota species, *Flavobacterium johnsoniae,* as a heterologous host, because *Runella* is not genetically tractable. To minimize interference from endogenous *F. johnsoniae* immune systems, we engineered a host strain lacking five defense loci: pAgo, Septu, Gabija, Tmn, and CBASS. We also sequenced and annotated two *F. johnsoniae* phages, φCj1 and φCj29, that were previously isolated but lacked genomic characterization^46^. Genome analysis revealed that φCj1 has a 352,212 bp genome encoding 600 protein-coding genes and 47 tRNAs, while φCj29 has a 240,799 bp genome with 338 protein-coding genes and no tRNAs, classifying them as a jumbo and a near-jumbo phage, respectively (Extended Data Fig. 9a).

Wild-type RzAgo provided strong protection against phage infection, increasing the plaque-forming threshold for φCj29 by over two orders of magnitude (Fig. 6a, b). Notably, protection against φCj1, which encodes its own set of tRNA genes, was weaker, suggesting that additional phage-encoded tRNAs may facilitate evasion of SLFN-mediated defense. This antiviral defense relies on both SLFN nuclease and ATP hydrolysis activity, as nuclease-deficient (D32A) and ATP hydrolysis-deficient (E314A) mutants each showed severely diminished protection (Fig. 6b). Interestingly, in human SLFN11, mutating nuclease pocket residue D252 enhances *in vitro* RNase activity^47^. When we mutated the conserved corresponding residue in RzAgo (D72A), protection against phage infection was strikingly enhanced relative to the wild-type (Fig. 6b). Together, these data highlight functional parallels between bacterial and mammalian SLFN domains and indicate that RzAgo employs nuclease activity of SLFN domain to prevent phage propagation.

**Figure 6.**
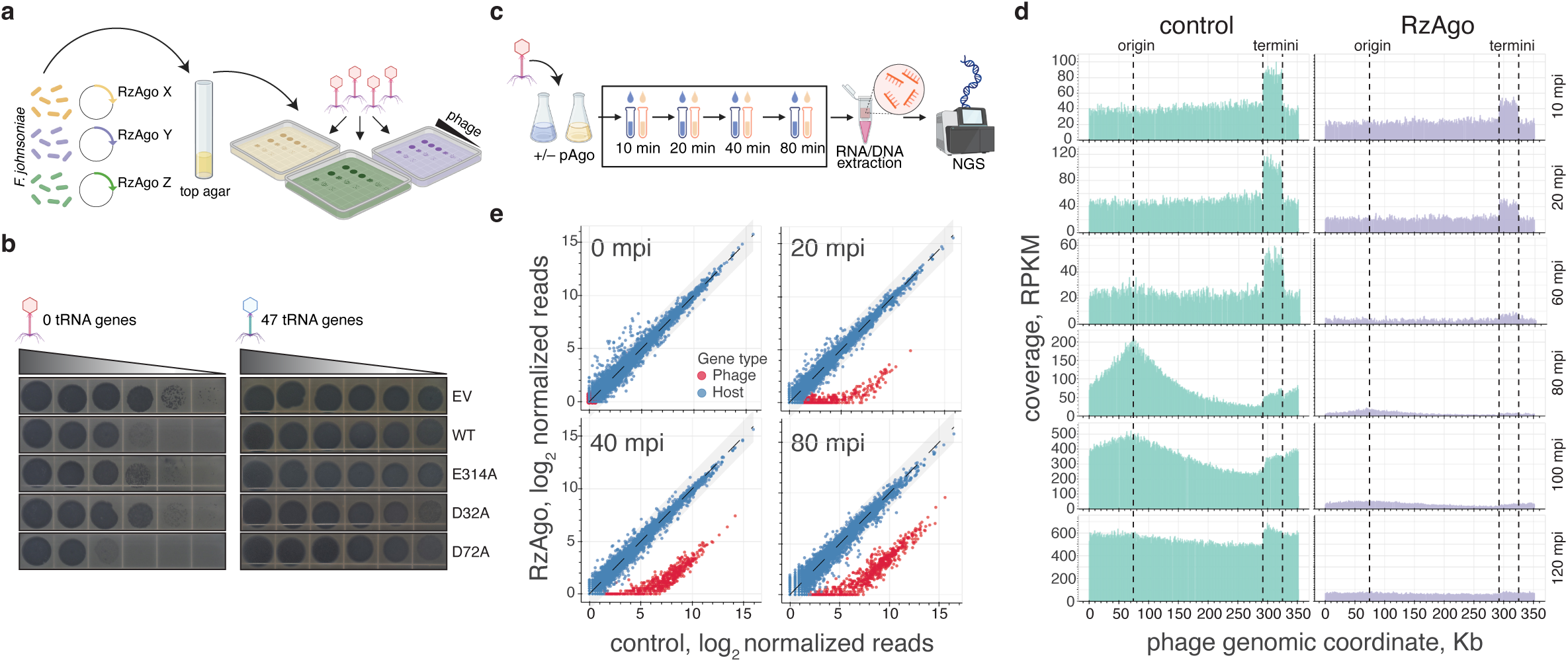
RzAgo protects cells from phage infection. **a**, Schematic of the efficiency-of-plating assay used to assess antiphage activity of RzAgo in *Flavobacterium johnsoniae*. **b**, Representative efficiency-of-plating assay for phages ϕCj29, left, and ϕCj1, right, on *F. johnsoniae* strains carrying an empty vector (EV), or expressing wild-type RzAgo (WT), the ATPase-deficient E314A mutant, the nuclease-deficient D32A mutant, or the hyperactive D72A mutant. **c**, Experimental strategy used to investigate the molecular basis of RzAgo-mediated phage defense. Cells expressing RzAgo or carrying an empty vector were infected with phage at a multiplicity of infection of approximately 5, and samples were collected at the indicated times after infection for DNA- and RNA-sequencing analyses. **d**, Genome-wide distribution of RPKM-normalized sequencing reads mapped to the ϕCj1 genome at successive time points after infection. Dashed lines mark the replication origin and long direct terminal repeats. **e**, Scatter plot of total RNA-seq data, shown as log₂ TMM-normalized reads, indicating gene expression changes during infection. Blue dots represent host genes and red dots represent phage genes. The shaded grey region denotes a ±1.5 log₂ fold-change interval; mpi – minutes post-infection.

To uncover the mechanism of RzAgo-mediated protection, we monitored infection progression via time-course RNA- and DNA-seq (Fig. 6c). In the absence of RzAgo, both phages proceeded through distinct early, middle, and late gene expression phases. Specifically, φCj1 genome circularization and replication only began following late gene expression, between 60 and 80 minutes post-infection (Fig. 6d, left; Extended Data Fig. 9b). Expression of the hyperactive RzAgo variant (D72A) severely disrupted this timeline. While φCj1 DNA was only moderately reduced (∼2-fold) at 20 minutes post-infection, it was largely eliminated by 60 minutes, directly blocking the onset of replication (Fig. 6d, right; Extended Data Fig. 9e, f). Concurrently, RzAgo induced a massive, ∼100-fold reduction in all φCj1 transcripts across all time points without significantly altering host transcript levels (Fig. 6e). A similar, though less pronounced, transcript depletion occurred during φCj29 infection (Extended Data Fig. 9c, d). This profound and specific destruction of phage RNA points to an abortive infection mechanism: target recognition unleashes SLFN-pAgo RNase activity, inducing host cell death to eliminate infected cells from the population and effectively prevent productive replication.

## DISCUSSION

Our structural and functional analyses establish SLFN-pAgos as a novel class of anti-phage defense systems that fuse a sequence-guided pAgo sensor and a SLFN nuclease effector into a single polypeptide, utilizing a GHKL domain for ATP-dependent tetramerization. In this system, the Ago core detects invading DNA, which in turn triggers the SLFN effector to cleave tRNA. We found that closely related SLFN domains have been exchanged between pAgo, NACHT and restriction-modification protein families (Fig. 1e, f). This architectural plasticity is widespread: Taboada et al. recently identified 55 distinct SLFN-containing domain configurations across bacteria and archaea, including an Ig-like sensor that detects a phage T5 protein^30^. Together, these modular arrangements point to a unifying evolutionary principle: conserved SLFN domains act as RNase effectors that are repeatedly paired with diverse sensory modules to detect distinct phage-derived signals. Although SLFN proteins were originally identified in mammals and long believed to be restricted to jawed vertebrates, they are now emerging as ancient components of cellular immunity. Given that the activation mechanisms of eukaryotic SLFNs remain poorly understood, the structural blueprint of SLFN-pAgos provides a critical foundation not only for understanding bacterial defense, but also for probing how SLFN proteins achieve target selectivity and activation in eukaryotes.

The structure of RzAgo tetramers provides a striking example of structural mimicry, revealing how bacterial and mammalian SLFNs achieve a shared functional geometry required for nuclease activity. In mammalian proteins such as human SLFN11, the SLFN domain comprises two topologically related subdomains: an N-lobe that provides a basic surface for nucleic acid binding, and a C-lobe carrying the nuclease active site^44^. Dimerization of these U-shaped domains creates a pincer-like asymmetric cleft that accommodates substrate nucleic acids^47,48^. Because SLFN-pAgos lack the N-lobe entirely, RzAgo overcomes this structural “deficit” through ATP-induced tetramerization. Within the tetramer, two SLFN domains dimerize to reconstitute the canonical bilobed geometry of a single extended mammalian SLFN. The subsequent association of two such dimers effectively recreates the mammalian substrate-binding cleft.

The transition of RzAgo to this functional state is driven predominantly by the GHKL domain, which mediates tetramer formation upon ATP binding. Similarly, ATP binding and hydrolysis drive large conformational rearrangements in other GHKL ATPases, such as Hsp90 and DNA gyrase B^49,50^. Remarkably, mammalian SLFN proteins also rely on an ATPase domain to drive active complex formation; however, the ATPase domains employed by these two systems are evolutionarily unrelated. SLFN-pAgos utilize a Bergerat-type GHKL fold, whereas mammalian SLFNs possess a distinct superfamily I (SF1) helicase/ATPase domain. Nucleotide-dependent switches that control effector oligomerization are widespread in innate immunity, including the assembly of supramolecular complexes by NACHT domains in both eukaryotic and prokaryotic NLR proteins^51,52^.

The use of nuclease effectors necessitates strict regulation to prevent spurious host toxicity. Our results suggest that SLFN-pAgos utilize oligomerization not only for activation but also for autoinhibition prior to target recognition. RzAgo–guide RNA complexes assemble into extended helical filaments, wherein the guide RNA acts as an intermolecular glue bridging adjacent protomers along a non-canonical trajectory. The head-to-tail packing of Argonaute core domains within the filament backbone physically forces the SLFN domains into spatially isolated clusters, a conformation fundamentally incompatible with the active tetrameric cleft. Filamentation thus locks the nuclease in a multimeric "safety catch". Target detection triggers a massive structural transition—unlike the localized shifts typical of eukaryotic Agos—disassembling the filament to release subunits competent for active tetramerization. This regulatory logic—utilizing a higher-order polymeric state to control effector domain activity—highlights a recurring theme in bacterial immunity. While CBASS and Thoeris utilize filamentation to activate their respective effectors^53,54^, RzAgo parallels Retron-Eco1, which cages its effector within an msDNA-stabilized filament to restrain toxicity^55^. Alongside Gabija and RADAR^56,57^, these systems illustrate that bacterial immune effectors frequently operate as dynamic supramolecular machines.

Together, our findings establish SLFN-pAgos as a paradigm for immune regulation via reversible structural switching. In this system, destructive nuclease activity is restrained within a guide-loaded filamentous state and unleashed upon sequence-specific target recognition coupled with ATP-driven tetramerization.

## ACKNOWLEDGEMENTS

We thank the Caltech cryo-EM facility, particularly Songye Chen, for expert assistance with cryo-EM data collection and Professors Po Lin Chiu and Ian MacRae for advice on cryo-EM data analysis. We are also grateful to Professor Mark McBride for his expert guidance on *Flavobacterium johnsoniae* and for kindly providing phage isolates used in this study. Special thanks to Denis Yudin for helpful discussions. This work was supported by the Merkin Translational Research grant to A.A.A., Caltech CEMI pilot grant to A.Kuzmenko and K.R., Caltech Baxter fellowship to A.Kuzmneko and Russian Science Foundation grant 22-14-00182-P to A.Kulbachinskiy.

## METHODS

### Strains and growth conditions

*Escherichia coli* NEB Turbo was used for routine plasmid propagation and molecular cloning. *E. coli* LOBSTR (DE3) (Kerafast) was used for recombinant protein expression.

*E. coli* BW25131 carrying pRK24 plasmid was as a donor stain for conjugation^58^. Plasmids were introduced into *E. coli* strains by chemical transformation using in-house prepared competent cells (Mix & Go *E. coli* Transformation Kit, Zymo Research) or by electroporation. *E. coli* strains were cultivated in LB medium at 37 °C or 30 °C (in case of the donor strain) supplemented with appropriate antibiotics where applicable (ampicillin, 100 μg ml^−1^; kanamycin, 50 μg ml^−1^; chloramphenicol, 12.5 μg ml^−1^; tetracycline, 10 μg ml^−1^). *Flavobacterium johnsoniae* DSM 6792 (DSMZ – German Collection of Microorganisms and Cell Cultures, Leibniz Institute) was cultivated in liquid or solid CYE medium (10 g l^−1^ casitone, 5 g l^−1^ yeast extract, 10 mM Tris-HCl pH 7.3, 8 mM MgSO_4_) at 30 °C with appropriate antibiotics where necessary (cefoxitin, 100 μg ml^−1^; erythromycin, 150 μg ml^−1^). Electrocompetent cells of *F. johnsoniae* were prepared from exponentially grown cultures by 3x washing with ice-cold 10 % glycerol. Plasmid was added to the cells in 1 cm cuvettes and electroporated at 400 Ω, 25 μF and 10 kV cm^-1^ (Gene Pulser II, Bio-Rad). *Runella zeae* DSM 19591 (DSMZ – German Collection of Microorganisms and Cell Cultures, Leibniz Institute) was grown on R2A agar plates at 30 °C.

Two bacteriophages infecting *F. johnsoniae* used in this study (φCj1 and φCj29) were a gift from Professor Mark McBride (University of Wisconsin-Madison) and were routinely propagated by the agarose overlay method^46^. Briefly, 200 μl of an overnight culture of *F. johnsoniae* was mixed with 10 μl of phage lysate to achieve an MOI of ∼0.1, and phage was allowed to adsorb for 30 minutes at room temperature without shaking. The phage–cell suspension was then mixed with 9 ml of molten 0.35% agarose in HTC medium (2 g l^−1^ tryptone, 0.5 g l^−1^ beef extract, 0.5 g l^−1^ yeast extract, 0.2 g l^−1^ sodium acetate, final pH 7.4) and poured over a solid HTC agar plate. After solidifying, plates were incubated overnight at 25 °C to achieve confluent lysis. The top agarose layer was scraped into 5 ml of SM buffer (50 mM Tris-HCl, 100 mM NaCl, 8 mM MgSO_4_, pH 7.4) and rocked gently overnight at 4 °C to allow phage particles to diffuse from the gel matrix. The lysate was centrifuged at 3,500g at 4 °C, passed through a 0.45 μm PES syringe filter, and stored at 4 °C. Phage titres were routinely verified by plaque assay, and phages were repurified by passaging through individual plaques after every three rounds of liquid propagation to prevent accumulation of unwanted mutations.

### Plasmid construction

The wild-type *RzAgo* gene was PCR-amplified from *R. zeae* genomic DNA using Q5 2× Master Mix (NEB) or repliQa HiFi ToughMix (Quantabio) and inserted into pET28b for heterologous expression in *E. coli* as an N-terminal His_8_-SUMO fusion by Gibson assembly (NEB). The construct was designed such that Ulp1p SUMO-protease cleavage yields a wild-type RzAgo protein carrying a single additional N-terminal glycine residue. All amino acid substitutions were introduced using the QuikChange site-directed mutagenesis kit (Agilent). Domain deletions were constructed by amplifying the plasmid with primers annealing to regions flanking the desired deletion, followed by circularization by Gibson assembly.

A replicative plasmid for expression of RzAgo in *F. johnsoniae* (pFj3_RzAgo) was constructed by PCR-amplifying the *RzAgo* gene together with its native promoter from *R. zeae* genomic DNA and inserting the fragment by Gibson assembly into the pCP29 backbone^59^, which was linearized by PCR to remove the *ErmF* gene (region between the AflII and BamHI sites). The resulting construct is a shuttle vector containing a high-copy-number pUC origin of replication and AmpR for propagation in *E. coli*, and a *Flavobacterium*-specific origin of replication and CfxA for propagation in *F. johnsoniae*. Mutant variants of RzAgo were subsequently re-cloned into the pFj3 backbone from their pET28b templates by PCR and Gibson assembly.

To introduce gene deletions into *F. johnsoniae*, we constructed a suicide vector (pFj2) by inserting the SacB negative-selection marker under the strong *F. johnsoniae* OmpA promoter into the pCP11 backbone, linearized by PCR between two PvuII sites, by Gibson assembly. For each target gene, ∼1.5 kb regions flanking the desired deletion were PCR-amplified and cloned into pFj2 by Gibson assembly. All final constructs were verified by Sanger (Laragen) or nanopore sequencing (Plasmidsaurus).

### Generation of *F. johnsoniae* knock-out strains

To minimize the contribution of native defence systems when assessing RzAgo antiviral activity, we constructed a strain of *F. johnsoniae* carrying deletions of five known defence loci — pAgo, tmn, Septu, Gabija, and gasdermin — identified using PADLOC (v1.1.0)^60^ with default parameters. This strain is hereafter referred to as FjΔ5DL. Sequential single deletions were introduced into the wild-type background using the pop-in/pop-out method. In each round, the relevant pFj2-based deletion vector was introduced into the recipient *F. johnsoniae* by conjugation from a donor *E. coli* strain. Briefly, equal volumes of exponentially grown donor and recipient cells (matched by OD_600_) were washed twice with fresh CYE medium, concentrated by resuspension in a small volume of residual medium after the final centrifugation step, mixed at an approximately 1:1 ratio, and spotted onto CYE agar. Conjugation was allowed to proceed overnight at 30 °C, after which cells were scraped into fresh CYE medium, serially diluted, and plated on CYE agar containing erythromycin to select for transconjugants (first recombination event). Counterselection against the *E. coli* donor was not required, as the *ErmF* gene cannot confer erythromycin resistance in *E. coli*. Individual transconjugants were re-streaked onto fresh selective plates after 2 days to eliminate any residual donor cells. To select for the second recombination event (pop-out), transconjugants were inoculated into liquid CYE medium, grown overnight at 30 °C, serially diluted, and plated on CYE agar supplemented with 5% sucrose for SacB-mediated counterselection. Successful deletions were identified by colony PCR and confirmed by Sanger sequencing.

### Sequencing, assembly and annotation of bacterial genomes

High-molecular-weight genomic DNA for nanopore sequencing was extracted from exponentially grown bacteria using the Monarch HMW DNA Extraction Kit for Tissue (NEB) following the manufacturer’s instructions for gram-negative bacteria. Nanopore long-read libraries were prepared and sequenced at the Millard and Muriel Jacobs Genetics and Genomics Laboratory (Caltech) on a MinION flow cell using R10.4 chemistry.

Hybrid genome assembly was performed using Unicycler (v0.4.8)^61^ with default parameters, combining the nanopore long reads with paired-end Illumina reads downloaded from the SRA (accession SRR896071 for *R. zeae* DSM 19591; SRR5188344 for *F. johnsoniae* DSM 6792) using the SRA Toolkit. Only reads with a quality score greater than 10 were used for assembly. Genome annotation was performed using the NCBI PGAP pipeline^62^ (Docker image 2022-04-14.build6021 for initial annotation; 2025-05-06.build7983 for final GenBank submission) with default parameters.

### Sequencing, assembly and annotation of bacteriophage genomes

Phage genomic DNA was extracted from high-titre lysate (≥10^10^ pfu ml^−1^) as described previously^63^. Briefly, 450 μl of lysate was treated with 2 μl Turbo DNase (Thermo Fisher) and 2 μl RNase A (DNase- and protease-free, Thermo Fisher) for 1 hour at 37 °C to remove bacterial nucleic acids. The reaction was quenched with EDTA (20 mM final concentration), and capsids were digested with 2.5 μl Proteinase K (molecular biology grade, NEB) for 1.5 hours at 56 °C. The lysate was mixed 1:1 (v/v) with buffer AL (DNeasy Blood & Tissue Kit, Qiagen) and incubated for 10 minutes at 70 °C, followed by addition of one volume of 95% ethanol. DNA was bound to a DNeasy Mini spin column, washed twice with buffers AW1 and AW2, and eluted in 30 μl buffer AE.

Phage genomic DNA was sheared by sonication using a Bioruptor Plus (Diagenode) to achieve a final fragment range of 200–600 bp. Sequencing libraries were prepared using the NEBNext Ultra II DNA Library Prep Kit for Illumina (NEB) according to the manufacturer’s instructions and sequenced on an Illumina NextSeq 2000 in 150 nt paired-end mode at the Millard and Muriel Jacobs Genetics and Genomics Laboratory (Caltech).

Paired-end reads were pre-processed using a custom adapter-trimming script to remove reads shorter than 14 nt and to filter reads with perfect alignment to the host genome. Phage genomes were assembled using SPAdes (v3.15.4)^64^ with options --careful and -k 21,33,55,77. Initial genome annotation was performed with Pharokka (v1.5.1)^65^ using default parameters. To improve annotation of unknown ORFs, structural information was incorporated: AlphaFold3^66^ models of phage proteins were generated using a standalone AlphaFold3 installation with default parameters, and these models were provided as input to Phold (v1.2.4)^67^, which uses Foldseek^68^ to identify structural homologs in the phage protein database.

### Protein expression and purification

Wild-type RzAgo and all mutant variants were expressed and purified following the same optimized protocol. The expression plasmid was introduced into *E. coli* LOBSTR (DE3) cells (Kerafast) by electroporation and grown on LB agar overnight at 37 °C. The following day, cells were scraped into 500 ml of autoinduction medium (AIM) and aerated at 37 °C until OD_600_ reached ∼1.0, after which cultures were transferred to 18 °C and grown for an additional 24 hours. Cells were harvested by centrifugation at 6,000g at 4 °C, and the pellet was resuspended in buffer A (50 mM HEPES-NaOH pH 7.6, 500 mM NaCl, 20 mM imidazole, 1 mM TCEP, 5% glycerol) at 5 ml per gram of wet cell weight. All subsequent steps were performed at 4 °C. Lysozyme was added to 1 mg ml^−1^ and the suspension was incubated for 30 minutes with gentle agitation. Complete EDTA-free protease inhibitors (Roche) were added, and cells were lysed by three passages through a high-pressure homogenizer at 30 kpsi (Constant Systems). Insoluble material was removed by centrifugation at 35,000g for 45 minutes.

The cleared lysate was applied to a 5 ml Ni-charged IMAC column (Bio-Rad) connected to a Bio-Rad NGC chromatography system and washed with 10 column volumes of buffer A. Co-purifying nucleotides were removed by washing with 10 column volumes of buffer A supplemented with 1 M NaCl, after which the protein was eluted with buffer A containing 300 mM imidazole. The eluate was transferred into buffer B (50 mM HEPES-NaOH pH 7.6, 300 mM NaCl, 10 mM imidazole, 1 mM TCEP, 5% glycerol) by on-column desalting (Cytiva) and treated overnight with in-house-prepared Ulp1 SUMO-protease (∼1:100 molar ratio). Precipitated material was removed by centrifugation at 12,000g for 15 minutes, and untagged RzAgo was isolated by reverse IMAC, passing the supernatant through a 1 ml Ni-charged HP IMAC column (Cytiva) at 0.5 ml min^−1^; cleaved, untagged RzAgo was collected in the flow-through.

To remove any residual nucleic acids co-purifying with RzAgo, the flow-through fraction was applied to a heparin column equilibrated with buffer C (20 mM HEPES-NaOH pH 7.6, 300 mM NaCl, 1 mM TCEP, 5% glycerol), washed until A_280_ reached baseline, washed further with 10% buffer D (buffer C supplemented with 1 M NaCl), and eluted in a step of 60% buffer D. Fractions containing RzAgo were pooled, concentrated using an Amicon ultrafiltration unit (50 kDa MWCO), and subjected to size-exclusion chromatography on a Superdex 200 Increase 10/300 GL column (Cytiva) into buffer E (40 mM HEPES-KOH pH 7.4, 250 mM potassium acetate, 8 mM magnesium acetate). Protein purity was assessed by SDS-PAGE, and successful removal of co-purifying nucleic acids was confirmed by an A_260_/A_280_ ratio below 0.6. Purified protein was concentrated to 50 μM, snap-frozen in single-use aliquots in liquid nitrogen, and stored at −80 °C.

### Production and purification of anti-RzAgo antibodies

Purified wild-type RzAgo was submitted for custom rabbit polyclonal antibody production (Albabion). The crude antibody-containing fraction was isolated from serum by ammonium sulfate precipitation and dissolved in PBS. Affinity purification was performed by applying the resuspended fraction to a custom column containing RzAgo covalently coupled to NHS-activated Sepharose 4 Fast Flow resin (Cytiva) at 0.1 ml min^−1^ at 4 °C. Bound antibodies were washed with 10 column volumes of PBS, followed by 5 column volumes of PBS supplemented with 500 mM NaCl, and eluted in 1 ml fractions with 100 mM glycine-HCl pH 3.0 directly into tubes containing 100 μl of 1 M Tris-HCl pH 8.5 for immediate neutralization. Fractions containing anti-RzAgo antibody were identified by SDS-PAGE, pooled, and buffer-exchanged into PBS supplemented with 10% glycerol by on-column desalting. The final antibody was concentrated using an Amicon ultrafiltration unit (30 kDa MWCO) to approximately 1 mg ml^−1^ and stored in single-use aliquots at −20 °C.

### *In vitro* nucleic acid binding assays

Binding affinities of wild-type RzAgo and the ATPase domain deletion mutant (SchlatpaseΔ; residues 1–442 deleted) towards RNA and DNA guides carrying different 5′-nucleotides were measured using a double-filter binding assay. Oligonucleotide substrates (5′-NUUAGACUUUAAGUCAAU-3′ for RNA; 5′-NTTAGACTTTAAGTCAAT-3′ for DNA) were 5′-radiolabeled by incubating 1 μM oligonucleotide with 1 μl of γ-^32^P-ATP (3,000 Ci mmol^−1^, Revvity) and polynucleotide kinase (PNK, NEB) in a 20 μl reaction. Twofold serial dilutions of protein (from 1,024 nM to 1 nM) were mixed with 50 nM radiolabeled guide in binding buffer (10 mM HEPES-NaOH pH 7.0, 100 mM NaCl, 5 mM MgCl_2_, 5% glycerol, 100 μg ml^−1^ BSA) and incubated for 30 minutes at 30 °C. Samples were filtered through a sandwich of nitrocellulose membrane (Cytiva; pre-activated by incubation in 0.4 M KOH for 10 minutes, washed extensively with ultrapure water, and equilibrated in binding buffer) over Hybond N+ membrane (Cytiva) mounted in a dot-blot apparatus (Bio-Rad). The sandwich was washed three times with binding buffer, air-dried for 15 minutes at 60 °C, exposed to a phosphor-storage screen (Cytiva) for 2 hours, and scanned on a Typhoon fluorescent imager (Cytiva).

Target nucleic acid binding was measured analogously. Binary complexes of RzAgo loaded with a 5′-A-phosphorylated, unlabeled RNA guide were first assembled at 2:1 protein:guide molar ratio for 15 minutes at 30 °C. Twofold serial dilutions of the binary complex (from 204.8 nM to 0.2 nM) were then mixed with 100 nM radiolabeled target (5′-UUUAUCAAAAAGAGUAUUGACUUAAAGUCUAACCUAUAGGAUACUUACAG-3′ for RNA target; 5′-TTTATCAAAAAGAGTATTGACTTAAAGTCTAACCTATAGGATACTTACAG-3′ for DNA target) and incubated for an additional 15 minutes before filtration as above. Raw signal was extracted using ImageQuant TL software (Cytiva) and apparent dissociation constants (K_d_) were determined by fitting to a single-site binding model in Prism 8 (GraphPad).

### ATP binding and hydrolysis

ATP binding was measured using the Differential Radial Capillary Action of Ligand Assay (DRaCALA)^69^. All binding assays were performed with apo protein in the absence of nucleic acid cofactors. Briefly, twofold serial dilutions of wild-type RzAgo or the ATPase-dead mutant (E314A) were mixed with α-^32^P-ATP (3,000 Ci mmol^−1^, Revvity) in a 20 μl reaction volume (0.5× buffer E, 100 μg ml^−1^ BSA) to a final concentration of ∼3.3 nM α-^32^P-ATP, and incubated at 30 °C for 2 minutes. No difference in binding was observed between 2- and 20-minute incubations; the shorter incubation was used to minimize ATP hydrolysis during the assay. A 1.5 μl aliquot of each reaction was spotted onto a 0.45 μm nitrocellulose membrane and allowed to absorb and dry. The membrane was exposed to a phosphor-storage screen (Cytiva) for 2 hours and scanned on a Typhoon fluorescent imager (Cytiva). Raw pixel values were extracted using ImageQuant TL software (Cytiva) and binding isotherms were fitted using a custom Python script. For nucleotide competition assays, unlabeled ribonucleoside triphosphates were added to a final concentration of 0.5 mM after 5 minutes of initial incubation with α-^32^P-ATP, and the reaction was continued for an additional 30 minutes before spotting.

ATP hydrolysis was assessed by thin-layer chromatography (TLC). All hydrolysis assays were performed with apo protein in the absence of nucleic acid cofactors. Wild-type RzAgo or the E314A mutant (500 nM) was mixed with γ-^32^P-ATP (3,000 Ci mmol^−1^, Revvity) in a 20 μl reaction volume (0.5× buffer E, 100 μg ml^−1^ BSA) to a final concentration of 4 nM γ-^32^P-ATP and incubated at 30 °C for 1 hour. Reactions (1.5 μl) were spotted onto PEI-cellulose TLC plates (Sigma) and developed in 1.5 M KH_2_PO_4_ (pH 3.8). Plates were dried, exposed to a phosphor-storage screen for 2 hours, and scanned on a Typhoon fluorescent imager.

### Analysis of protein complexes by crosslinking SDS-PAGE

Binary complexes of wild-type RzAgo or the indicated mutants (2 μM) were assembled by incubation with a 5′-phosphorylated 18 nt RNA guide (5′-AUUAAGAGAUGUUGAUGA-3′; 4 μM) in 0.5× buffer E supplemented with 500 μM of ATP or γ-S-ATP for 15 minutes at 37 °C. Ternary complexes were subsequently assembled by addition of a complementary 24 nt target DNA (5′-AACTCATCAACATCTCTTAATTTA-3′; 8 μM) and incubation for a further 20 minutes at 37 °C. Apo-form control samples were prepared identically, substituting RNA and DNA additions with an equivalent volume of nuclease-free water. All samples were normalized with respect to buffer composition before crosslinking.

Glutaraldehyde was added to a final concentration of 0.025% (v/v) — the minimum concentration that preserved quaternary structure without introducing over-crosslinking artefacts, as determined in a series of preliminary experiments — and crosslinking was allowed to proceed for 10 minutes at room temperature. Reactions were quenched with 40 mM Tris-HCl (pH 7.4) for 15 minutes at room temperature. Samples were resolved on 3–8% Tris-acetate gels (Thermo Fisher) and stained overnight with One-Step Blue protein gel stain (Biotium). Gels were imaged on a Li-COR Odyssey infrared imager.

### *In vitro* tRNA cleavage assay

To assess the nuclease activity of RzAgo *in vitro*, apo (no guide or target), binary (RNA guide-loaded), and ternary (RNA guide- and DNA target-loaded) complexes were assembled and challenged with total *E. coli* tRNAs (Fermatix). Briefly, 2 μM RzAgo was incubated with a 1.25-fold molar excess of a 5′-phosphorylated 18 nt RNA guide for 20 minutes at 37 °C, followed by addition of a 1.5-fold molar excess (relative to protein) of either a complementary or a non-complementary (5′-AACCATTCATTACCAACAATTCCA-3′) 24 nt DNA target and incubation for a further 15 minutes. Apo-form control samples were prepared identically, substituting guide and target additions with equivalent volumes of nuclease-free water. All samples were normalized to identical buffer conditions (20 mM HEPES-KOH pH 7.5, 100 mM potassium acetate, 5 mM magnesium acetate, 5 mM manganese acetate, 5 mM ATP) before initiating the reaction.

Cleavage reactions were initiated by addition of total *E. coli* tRNA to a final concentration of 1 μM. At each indicated time point, an aliquot was withdrawn and quenched by addition of an equal volume of stop solution (8 M urea, 20 mM EDTA). Proteinase K was then added separately (0.5 μl of 20 mg ml⁻¹ stock (NEB)) and samples were incubated at 50 °C for 15 minutes to ensure complete protein digestion before gel loading. Samples were resolved on a 15% denaturing TBE–urea polyacrylamide gel at 250 V for 55 minutes. Gels were stained with SYBR Gold, scanned on a Typhoon fluorescent imager (Cytiva), and raw images were processed in Fiji.

### Isolation of RzAgo-associated small RNAs from bacteria

To isolate RzAgo-associated small RNAs from *R. zeae*, cells were spread from a frozen seed stock onto 120 × 120 mm R2A agar plates (∼100 μl per plate at OD_600_ ∼1) and grown at 30 °C for 24 hours. Cells were scraped into ice-cold immunoprecipitation (IP) buffer (25 mM HEPES-KOH pH 7.05, 150 mM potassium acetate, 5 mM magnesium acetate, 0.5% CHAPS) supplemented with RNaseOUT recombinant RNase inhibitor (Thermo Fisher) and the suspension was adjusted to OD_600_ ∼5. All subsequent steps were performed at 4 °C. Cells were lysed by indirect sonication in a Bioruptor Plus (Diagenode) for 15 cycles (30 s on/30 s off, high amplitude), and insoluble material was removed by centrifugation at 21,000g for 15 minutes. Total protein concentration was measured using the Pierce Detergent-Compatible Bradford Assay (Thermo Fisher) and the lysate was normalized to 1 mg ml^−1^. RzAgo–nucleic acid complexes were immunoprecipitated by adding 5 μl of anti-RzAgo rabbit polyclonal antibody (∼1 mg ml^−1^) and rocking for 4 hours, followed by 2 hours of incubation with 20 μl of protein A/G magnetic beads (Thermo Fisher). Beads were washed five times with IP buffer (5 minutes each) and resuspended in 300 μl of elution buffer (100 mM Tris-HCl pH 7.5, 150 mM NaCl, 1 mM EDTA, 1% SDS).

The sample was digested with 1 μl Proteinase K (NEB) for 1 hour at 56 °C. Nucleic acids were extracted using phenol:chloroform:isoamyl alcohol (25:24:1, pH 8.0, Sigma) with a phase-lock tube (MaXtract High-Density, Qiagen) and precipitated with ethanol in the presence of GlycoBlue coprecipitant (Thermo Fisher). The pellet was dissolved in 25 μl ultrapure water.

RzAgo-associated small RNAs were size-selected by denaturing PAGE. For visualization, a 5 μl aliquot was dephosphorylated with rSAP (NEB) and 5′-radiolabeled with γ-^32^P-ATP using PNK (NEB). The labeled aliquot was combined with the unlabeled sample, mixed with an equal volume of denaturing sample buffer (8 M urea, 20 mM EDTA, 0.005% bromophenol blue, 0.005% xylene cyanol), and resolved on a 19% polyacrylamide–urea gel under denaturing conditions. Gel slices corresponding to 14–25 nt RNAs were excised, crushed, and RNA was eluted overnight in 0.4 M NaCl at 21 °C with constant agitation. RNA was recovered by ethanol precipitation and resuspended in 20 μl ultrapure water.

Small RNA sequencing libraries incorporating unique molecular identifiers (UMIs) were prepared according to^70^ with the following modifications. The 3′-adapter (5′-*NNNgtcNNNtagNNN*AGATCGGAAGAGCACACGTCT-3′, UMI positions in italic) and 5′-adapter (5′-GTTCAGAGTTCTACAGTCCGACGATC*NNNcgaNNNtacNNN*-3′, UMI positions in italic) were redesigned for compatibility with the NEBNext small RNA sequencing workflow. cDNA synthesis was performed using the NEBNext SR RT primer (5′-AGACGTGTGCTCTTCCGATCT-3′). Final PCR indexing was performed using KAPA HiFi HotStart ReadyMix (Roche) with NEBNext small RNA indexing primers; cycle numbers were optimized empirically to avoid overamplification. Libraries were gel-purified on a 6% TBE gel (Thermo Fisher), stained with SYBR Gold, excised, and eluted overnight in 0.4 M NaCl at 21 °C with agitation. After ethanol precipitation, libraries were resuspended in 10 mM Tris-HCl pH 8.5 and quantified using the Qubit dsDNA HS Assay Kit (Thermo Fisher). Libraries were sequenced on an Illumina NextSeq 2000 in 100 nt single-end read mode at the Millard and Muriel Jacobs Genetics and Genomics Laboratory (Caltech).

### Phage plaque assay

*F. johnsoniae* FjΔ5DL strains expressing the indicated RzAgo variants from a pFj3 plasmid under its native promoter were grown overnight at 30 °C with shaking at 200 rpm in CYE medium supplemented with 100 μg ml^−1^ cefoxitin. For each assay, 200 μl of overnight culture was mixed with 9 ml of molten 0.35% agarose in HTC medium supplemented with cefoxitin, poured over an HTC agar plate, and allowed to solidify and dry at room temperature for 30 minutes. Tenfold serial dilutions of phage were spotted on the overlay, allowed to dry, and plates were incubated overnight at 25 °C. Plaque images were acquired using a custom photo box equipped with an Olympus camera against a black background.

### Time course analysis of phage infection

*F. johnsoniae* FjΔ5DL carrying pFj3_RzAgoD72A or the empty vector control were grown in CYE medium to OD_600_ ∼1 and infected with φCj1 or φCj29 at an MOI of ∼5. Infections were carried out at 25 °C without shaking. After 15 minutes, cultures were washed with fresh CYE medium to remove unabsorbed phage particles (3,200g for 1 minute) followed by resuspension in fresh CYE. The t = 0 sample was collected before addition of the phage. Duplicate 1 ml samples (one for RNA-seq and one for DNA-seq) were removed at the indicated time points, centrifuged at 6,000g for 30 seconds, and the pellets were immediately frozen in liquid nitrogen and stored at −80 °C until processing.

Total RNA was isolated using the RNAsnap method^71^. Briefly, pellets were resuspended in 100 μl of RNA extraction solution (18 mM EDTA, 0.025% SDS, 1% 2-mercaptoethanol, 95% formamide) and boiled at 95 °C for 7 minutes. Samples were centrifuged at 16,000g for 15 minutes, and the supernatant was transferred to a new tube. Samples were diluted 1:4 with ultrapure water and RNA was precipitated with ethanol. Crude RNA was purified using an RNA Clean and Concentrator column (Zymo Research) with on-column DNase I treatment according to the manufacturer’s instructions. RNA concentration was quantified using the Qubit RNA Broad Range Assay Kit (Invitrogen). Ribosomal RNA was removed using the Pan-Prokaryote riboPOOL rRNA depletion kit as per manufacturer’s guidelines. RNA-seq libraries from φCj29-infected samples were prepared using the NEBNext Ultra II Directional RNA Library Prep Kit (NEB). RNA-seq libraries from φCj1-infected samples were prepared using the TGIRT-seq workflow as described^72^.

Total DNA was extracted from frozen pellets using the Monarch Spin gDNA Extraction Kit (NEB) following the manufacturer’s instructions. DNA-seq libraries were prepared using the NEBNext Ultra II DNA Library Prep Kit for Illumina (NEB). All libraries were sequenced on an Illumina NextSeq 2000 in 100 nt single-end read mode at the Millard and Muriel Jacobs Genetics and Genomics Laboratory (Caltech).

### Analysis of high-throughput sequencing data

#### Analysis of RzAgo-associated small RNAs

Adapter sequences (5′-AGATCGGAAGAGCACACGTCTGAACTCCAGTCAC-3′) and reads shorter than 14 nt were removed using cutadapt (v4.2). UMIs were extracted from trimmed reads using UMI-tools (v1.1.4)^73^ with default parameters, and read quality was assessed using FastQC (v0.11.9). UMI-extracted reads were mapped to the *R. zeae* DSM 19591 genome in two passes using bowtie (v1.3.1): uniquely mapping reads were first identified with parameters -v 0 -m 1, and multimapping reads were subsequently realigned with parameters -a --best --strata -v 0 -m 10000. Only uniquely mapped reads were retained for downstream analysis. Aligned reads were converted to BAM format, sorted, and indexed using samtools (v1.16.1), then deduplicated based on UMI sequences using the UMI-tools dedup module with default parameters. Read length distributions were calculated and plotted using a custom R script. Nucleotide composition logos were generated from deduplicated, uniquely mapped reads trimmed to 14 nt using SeqKit (v2.12.0) and visualized using WebLogo3 (v3.9.0).

#### Analysis of total RNA-seq during phage infection

For φCj29-infected samples, adapter sequences were removed and reads shorter than 16 nt were discarded using cutadapt. Reads were mapped to both the *F. johnsoniae* DSM 6792 and phage genome sequences with up to 2 mismatches allowed using bowtie, retaining only uniquely mapped reads. For φCj1-infected samples, adapter trimming and minimum length filtering were performed identically with cutadapt, after which UMIs were extracted using UMI-tools for downstream positional deduplication tracking; reads were then mapped to the host and phage genomes using bowtie2 (--very-sensitive-local). For both libraries, gene-level count matrices were generated using featureCounts (v2.0.6) with default parameters and normalized using the TMM method implemented in the RNAnorm Python package (v2.1.0). Coverage plots and other visualizations were generated using custom Python scripts.

#### Analysis of total DNA-seq during phage infection

Adapter sequences and reads shorter than 14 nt were removed using cutadapt, and read quality was assessed using FastQC. Reads were mapped to both the *F. johnsoniae* DSM 6792 and phage genome sequences with up to 1 mismatch allowed using bowtie, retaining only uniquely mapped reads. Aligned reads were sorted and indexed using samtools, and genome-wide coverage was calculated using the bamCoverage function of deepTools (v3.5.1) with RPKM normalization (--normalizeUsing RPKM). Coverage tracks were visualized using custom Python script.

### Phylogenetic analysis

#### Phylogeny of prokaryotic Argonaute proteins

Sequences of prokaryotic Argonaute (pAgo) proteins were searched in the NCBI RefSeq protein database using deltaBLAST with the conserved MID and PIWI domain sequences from five previously characterized pAgos as queries (TtAgo: WP_011229221.1, CbAgo: WP_045143632.1, KmAgo: WP_010289662.1, MpAgo: WP_014295921.1, RsAgo: WP_011910606.1). Identical sequences were removed using MMseqs2 hash-based clustering (mmseqs clusthash), and sequence diversity was further reduced by clustering at 80% identity over 80% coverage using MMseqs2. For multiple sequence alignment (MSA) generation, MID and PIWI domain sequences were extracted from cluster representatives and were clustered further at 70% sequence identity. The MSA was generated using MAFFT with the L-INS-i strategy and trimmed using trimAl with a gap threshold of 0.3 (-gt 0.3). A maximum-likelihood phylogenetic tree was constructed from this alignment using IQ-TREE (v3.0.1) with 1,000 ultrafast bootstrap replicates and 1,000 SH-aLRT replicates (-B 1000 -alrt 1000 -bnni), with the best-fit partitioned substitution model selected automatically using ModelFinder (-m MFP+merge), yielding Q.PFAM+F+R10 for the MID domain partition and LG+F+R10 for the PIWI domain partition. The resulting tree was visualized and annotated in iTOL.

#### Phylogeny of Slfn-pAgo proteins

Sequences of Slfn-pAgo fusion proteins were identified by running deltaBLAST against the NCBI non-redundant (nr) protein database. Identical sequences were removed using MMseqs2 hash-based clustering, and a full-length MSA was constructed from the remaining sequences using MAFFT with the L-INS-i strategy. The corresponding species tree was generated using GTDB-Tk. To assess the extent of horizontal gene transfer (HGT) of Slfn-pAgo loci, the protein tree and species tree were compared by co-visualization as a tanglegram using a custom R script.

#### Structural phylogeny of prokaryotic Alba2-containing proteins

Prokaryotic Alba2-containing proteins were identified in the NCBI nr database using the Pfam profile PF04326. Sequence redundancy was first reduced using MMseqs2 hash-based clustering, and the remaining sequences were further clustered using Foldseek based on ProstT5-derived 3Di structural alphabet representations. Cluster representatives — covering approximately 3,400 structurally diverse proteins from an initial set of ∼115,000 hits — were modelled with AlphaFold3 using default parameters, and predicted structures were annotated using DPAM against the ECOD database. Alba2 domains were excised from the AlphaFold3 models and used to generate structure-derived 3Di sequences with Foldseek. These 3Di sequences were aligned using MAFFT with the local-pair iterative refinement strategy (L-INS-i) and the Foldseek 3Di substitution matrix supplied as a user-defined amino-acid substitution matrix. The resulting 3Di alignment was trimmed using trimAl (-gt 0.3), and a maximum-likelihood structural phylogenetic tree was built using IQ-TREE (v3.0.1) with 1,000 ultrafast bootstrap replicates and 1,000 SH-aLRT replicates (-B 1000 -alrt 1000 -bnni). The tree was visualized and annotated in iTOL.

### Negative stain electron microscopy

A 1.5% (w/v) uranyl formate solution was prepared by dissolving in boiling double-distilled water, filtered to remove large particles, aliquoted into 100 μl portions, and flash-frozen in liquid nitrogen. Carbon support film grids (300 mesh copper, Ted Pella no. 01843) were glow-discharged for 1 minute at 15 mA. RzAgo samples in the apo, binary complex (18 nt guide RNA), and ternary complex (18 nt guide RNA and 24 nt target ssDNA) states were prepared in 0.5× buffer E without ATP or ATP analogues. Each sample was prepared at 2 μM RzAgo with a 1.5x molar excess of guide RNA and 2x molar excess of target DNA where applicable, and incubated at room temperature for 30 minutes. Samples were crosslinked with 0.025% glutaraldehyde for 15 minutes and quenched with 40 mM Tris-HCl for 15 minutes at room temperature, then diluted to 200 nM RzAgo monomer. A 3 μl aliquot was applied to the carbon face of a glow-discharged grid and incubated for 1 minute. Excess sample was removed by blotting with Whatman filter paper, and the grid was stained twice with 3 μl uranyl formate (1 minute per application), blotting between applications. Grids were air-dried before imaging. Data were collected on a Talos Arctica (Thermo Fisher) at 200 kV equipped with a 6k x 4k Gatan K3 direct electron detector. A total of 1,800, 3,663 and 1,800 micrographs were collected for the apo, binary complex, and ternary complex datasets, respectively, at ×45,000 magnification with a total exposure dose of 65 e^−^ Å^−2^ and a pixel size of 0.869 Å. Data were imported into cryoSPARC^74^ and motion-corrected before particles were selected using Blob Picker or manual picking for 2D class averaging.

### Cryo-EM grid preparation, data collection, model building and refinement

#### Cryo-EM structure of the RzAgo apo monomer

Purified RzAgo was prepared at 12.5 μM in 0.5× buffer E without ATP. Quantifoil R 1.2/1.3 Au 400 grids were glow-discharged for 30 seconds at 15 mA (Pelco EasiGlow). A 4 μl aliquot was applied in a Vitrobot Mark IV (Thermo Fisher) at 25 °C and 100% humidity, blotted for 3 s at blot force 0 with no wait time, and plunge-frozen in liquid ethane. Data were collected on a Titan Krios (Thermo Fisher) at 300 kV equipped with a Gatan K3 6k x 4k direct electron detector and Gatan BioQuantum energy filter at ×130,000 magnification with a total exposure dose of 60 e^−^ Å^−2^; 3,256 movies were collected with 40 frames per movie. All data processing was performed in CryoSPARC. Micrographs were motion corrected using the Patch Motion correction job and the CTF was corrected with CTFFIND4. 3,108 exposures were accepted. Particles were picked with Blob Picker, inspected, extracted, and assembled into 2-dimensional class averages iteratively until an appropriate box size was established. Ab initio 3D classification (2 classes) and heterogeneous refinement (2 classes) were performed and the particle stack from the best class was used as training data for template picking using Topaz^75^. Five iterative rounds of Topaz particle picking, particle extraction, 2D and 3D classification, particle class rebalancing, and refinement were necessary. The final round of Topaz training yielded 578,612 unique particles, extracted with a box size of 320 pixels downsampled to 160 pixels. Particles were further sorted with heterogeneous refinement using 5 query volumes previously obtained, then non-uniform refinement of the best class and particle reextraction of the 219,569 selected particles to a box size of 320 pixels with no downsampling. A final non-uniform refinement yielded a map at 3.58 Å resolution. An AlphaFold3^66^ model of RzAgo was fitted into the density in ChimeraX^76^ (v.1.9) using ISOLDE^77^ and iteratively refined by real-space refinement in Phenix^78,79^ (v.1.21.2-5419-000) and manual correction in Coot^80,81^ (v.0.9.8.96), with final refinement with the EMAN2 package e2gmm^82^ prior to deposition.

#### Cryo-EM structure of the RzAgo apo tetramer

Purified RzAgo was prepared at 2 μM in 0.5× buffer E supplemented with γ-S-ATP. Samples were crosslinked with 0.025% glutaraldehyde for 15 minutes and quenched with 40 mM Tris-HCl for 15 minutes at room temperature. Quantifoil R 1.2/1.3 Cu 300 grids were glow-discharged for 30 seconds at 15 mA (Pelco EasiGlow). A 4 μl aliquot was applied in a Vitrobot Mark IV (Thermo Fisher) at 25 °C and 100% humidity, blotted for 4.5 s at blot force 0 with a 1-minute wait time, and plunge-frozen in liquid ethane. Data were collected on a Titan Krios (Thermo Fisher) at 300 kV equipped with a Gatan K3 6k x 4k direct electron detector and Gatan BioQuantum energy filter at ×130,000 magnification with a total exposure dose of 40 e^−^ Å^−2^; 6,451 movies were collected with 78 frames per movie. All data processing was performed in CryoSPARC. Micrographs were motion corrected using the Patch Motion correction job and the CTF was corrected with CTFFIND4. 5,374 exposures were accepted. 2,909,653 were picked with Blob Picker, inspected, and extracted with a box size of 600 pixels, resampled to 300 pixels. Particles were classified into 300 2D class averages, of which 28 were selected (322,600 particles). These were classified into 3 ab initio maps with C1 symmetry, and particles from the best class were passed through a round of homogeneous of refinement (109,024). This map was further refined with a second round of 2D classification and particle selection, and the final map was generated via non-uniform refinement. The final map has a reported resolution of 3.4 Å (GSFSC) and contained 64,577 particles. An AlphaFold3 model was fitted in ChimeraX (v.1.9) using ISOLDE and iteratively refined in Phenix (v.1.21.2-5419-000) and Coot (v.0.9.8.96). Ligand restraints were generated from SMILES strings using eLBOW (Phenix) and ligands were fitted manually in ISOLDE before iterative refinement, with final refinement with the EMAN2 package e2gmm prior to deposition.

#### Cryo-EM structure of the RzAgo binary complex dimer

Purified RzAgo was prepared at 2 μM in 0.5× buffer E (without ATP or γ-S-ATP), and the binary complex was assembled by addition of guide RNA at a 1.5-fold molar excess. Quantifoil R 1.2/1.3 Au 400 grids were glow-discharged for 30 seconds at 15 mA (Pelco EasiGlow). A 4 μl aliquot was applied in a Vitrobot Mark IV (Thermo Fisher) at 25 °C and 100% humidity, blotted for 3 s at blot force 0 with a 15 s wait time, and plunge-frozen in liquid ethane. Data were collected at 30° stage tilt on a Titan Krios (Thermo Fisher) at 300 kV equipped with a Gatan K3 6k x 4k direct electron detector and Gatan BioQuantum energy filter at ×105,000 magnification (pixel size 0.42 Å) with a total exposure dose of 60 e^−^ Å^−2^; 2,908 movies were collected. All data processing was performed in CryoSPARC. Micrographs were motion corrected using the Patch Motion correction job and the CTF was corrected with CTFFIND4, and 1,873 exposures were accepted. Using a subset of 324 micrographs, 138,848 particles were picked using the Blob picker with a particle diameter range of 100-300 Å, inspected, and extracted with a box size of 800 pixels, downsampled to 200. These were passed through two rounds of 2D classification and class rebalancing, yielding 16,653 accepted particles. These particles were used as a training data set for Topaz particle picking. Three rounds of Topaz training, opening to all accepted micrographs in the second round, followed by 2D classification, 3D ab initio map generation and heterogeneous refinement, were used to curate particles. A final stack of 85,382 particles extracted with a box size of 800 pixels was used to obtain a final structure at 4.03 Å via non-uniform refinement. An AlphaFold3 model of the RzAgo–guide RNA binary complex was fitted in ChimeraX (v.1.9) using ISOLDE and refined in Phenix (v.1.21.2-5419-000), with final refinement with the EMAN2 package e2gmm prior to deposition.

#### Cryo-EM structure of the RzAgo binary complex pentamer

Purified RzAgo was prepared at 4 μM in 0.5× buffer E supplemented with γ-S-ATP and magnesium acetate. Guide RNA was added and the sample was crosslinked with 0.025% glutaraldehyde for 15 minutes and quenched with 40 mM Tris-HCl for 15 minutes at room temperature. Quantifoil R 1.2/1.3 Cu 300 grids were glow-discharged for 30 seconds at 15 mA (Pelco EasiGlow). A 4 μl aliquot was applied in a Vitrobot Mark IV (Thermo Fisher) at 25 °C and 100% humidity, blotted for 4.5 s at blot force 0 with a 1-minute wait time, and plunge-frozen in liquid ethane. Data were collected on a Titan Krios (Thermo Fisher) at 300 kV equipped with a Gatan K3 6k x 4k direct electron detector and Gatan BioQuantum energy filter at ×130,000 magnification (pixel size 0.32 Å) with a total exposure dose of 45 e^−^ Å^−2^; 6,489 movies were collected. All data processing was performed in CryoSPARC. Micrographs were motion corrected using the Patch Motion correction job and the CTF was corrected with CTFFIND4. 6,165 micrographs were accepted via manual exposure curation. 2.9 million particles were initially picked using Blob picker with an expected particle diameter of 70-600 Å. Two rounds of 2D classification selected 232,476 particles, which were used for ab initio 3D map generation (3 maps requested). Each ab initio map was then refined using heterogeneous refinement, further selecting a promising particle population. Particle selection was further refined through three rounds of Topaz model training, which entailed training, extraction, one or more rounds of 2D classification, heterogeneous refinement, and then non-uniform refinement of the best class. A final particle stack containing 111,503 particles was extracted with a box size of 1024 pixels. Non-uniform refinement of 111,503 particles resulted in a final structure at 4.33 Å resolution. Model building used an AlphaFold3 model of RzAgo combined with the previously determined apo tetramer Slfn/ATPase domains and binary complex pAgo dimer as starting models. Initial fitting was performed in ChimeraX (v.1.9) using ISOLDE; iterative refinement was performed in Phenix (v.1.21.2-5419-000), with ligand restraints recycled from the apo tetramer refinement, with final refinement with the EMAN2 package e2gmm prior to deposition.

#### Cryo-EM structure of the RzAgo ternary complex

Purified RzAgo was prepared at 2 μM in 0.5× buffer E supplemented with 1 mM ATP. Guide RNA and target DNA were added to final concentrations of 1.5 μM and 1 μM, respectively. Quantifoil R 1.2/1.3 Cu 300 grids were glow-discharged for 30 seconds at 15 mA (Pelco EasiGlow). A 3.5 μl aliquot was applied in a Vitrobot Mark IV (Thermo Fisher) at 25 °C and 100% humidity, blotted for 3 s at blot force 0 with a 15 s wait time, and plunge-frozen in liquid ethane. Data were collected on a Titan Krios (Thermo Fisher) at 300 kV equipped with a Gatan K3 6k x 4k direct electron detector and Gatan BioQuantum energy filter at ×130,000 magnification (pixel size 0.338 Å) with a total exposure dose of 35 e^−^ Å^−2^; 12,916 movies were collected. All data processing was performed in CryoSPARC. Micrographs were motion corrected using the Patch Motion correction job and the CTF was corrected with CTFFIND4. 11,692 micrographs were accepted via manual exposure curation. 3.2 million particles were initially picked from a subset of 2,001 micrographs using Blob picker with an expected particle diameter of 50-200 Å, and 801,701 particles were extracted with a box size of 512 pixels (downsampled to 128). 2D classification selected 77,831 particles, which were used for ab initio 3D map generation (3 maps requested). Each ab initio map was then refined using heterogeneous refinement and the best refined by homogeneous refinement (31,884 particles). Further 2D classification from this refinement selected 28,114 particles, which were rebalanced and used for Topaz training on the 2,001-micrograph data set, which entailed training, extraction, one or more rounds of 2D classification, heterogeneous refinement, and then non-uniform refinement of the best class. Three rounds of Topaz training were performed, expanding the number of micrographs to the total data set on the second round. A final particle stack containing 122,907 particles was extracted with a box size of 512 pixels. After 2D classification, and heterogeneous refinement, non-uniform refinement of 76,858 particles resulted in a final structure at 3.28 Å resolution. An AlphaFold3 model was fitted in ChimeraX (v.1.9) using ISOLDE and refined in Phenix (v.1.21.2-5419-000), with final refinement with the EMAN2 package e2gmm prior to deposition.

## Data availability

Atomic coordinates and cryo-EM maps for all structures reported in this study have been deposited in the Protein Data Bank (PDB) and Electron Microscopy Data Bank (EMDB) under the accession codes: 35YR/EMD-77286 (RzAgo apo monomer), 35YS/EMD-77287 (RzAgo binary complex dimer), 35ZE/EMD-77296 (RzAgo binary complex pentamer), 35YQ/EMD-77285 (RzAgo ternary complex), and 35YP/EMD-77284 (RzAgo apo tetramer in complex with ATPγS). Raw sequencing data from time-course total RNA-seq, DNA-seq, and small RNA immunoprecipitation experiments have been deposited in the NCBI Sequence Read Archive (SRA) under BioProject accession PRJNA1470036. Assembled bacterial and phage genome sequences have been deposited in the NCBI Genome database under BioProject accession PRJNA1457876. All other data supporting the findings of this study are available from the corresponding authors upon reasonable request.

---

**Extended Data Figure 1.**
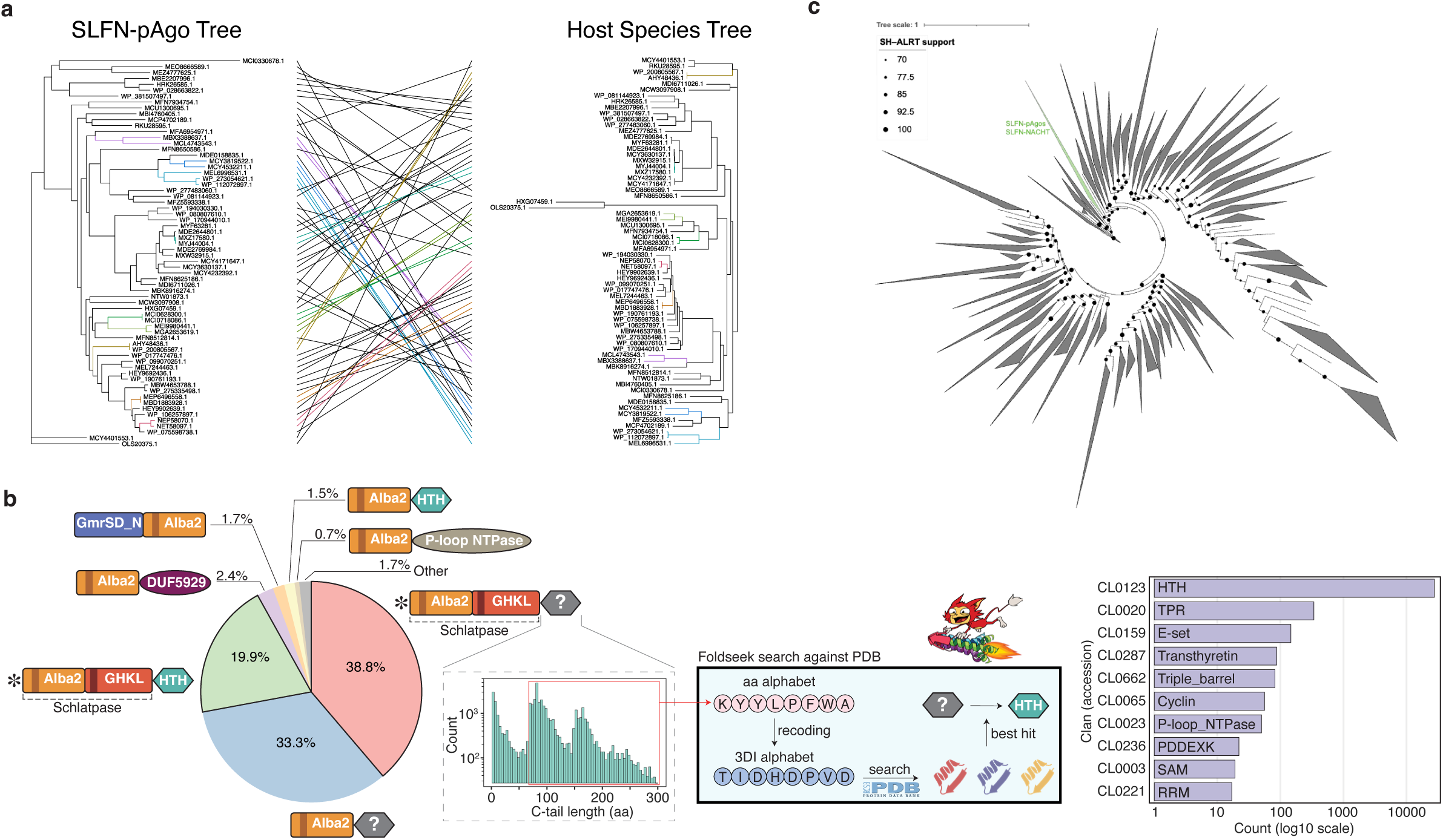
Bioinformatic characterization of SLFN-pAgos and other prokaryotic SLFN domain-containing proteins. **a,** Tanglegram illustrating incongruence between the evolutionary history of SLFN-pAgo proteins and that of their host species. The two trees show substantial topological discordance, with a Robinson–Foulds distance of 104/126 (82.5%). Only 66 SLFN-pAgos with robust genome annotations were included. **b,** Workflow and results of structural domain annotation for the C-terminal tails of Schlatpase-containing proteins. Many proteins contain additional domains not detected by sequence-based annotation tools. More than 90% of these proteins contain a helix–turn–helix (HTH) domain, followed by fusions to TPR and Ig-like domains. **c,** Maximum-likelihood structural phylogeny of prokaryotic SLFN_alba2 domains. The tree was constructed using AlphaFold3-predicted SLFN_alba2 domain structures from 3,394 structural cluster representatives (see Methods). The clade containing SLFN-pAgo and SLFN-NACHT proteins is highlighted in green. Dots indicate SH-aLRT branch support values.

**Extended Data Figure 2.**
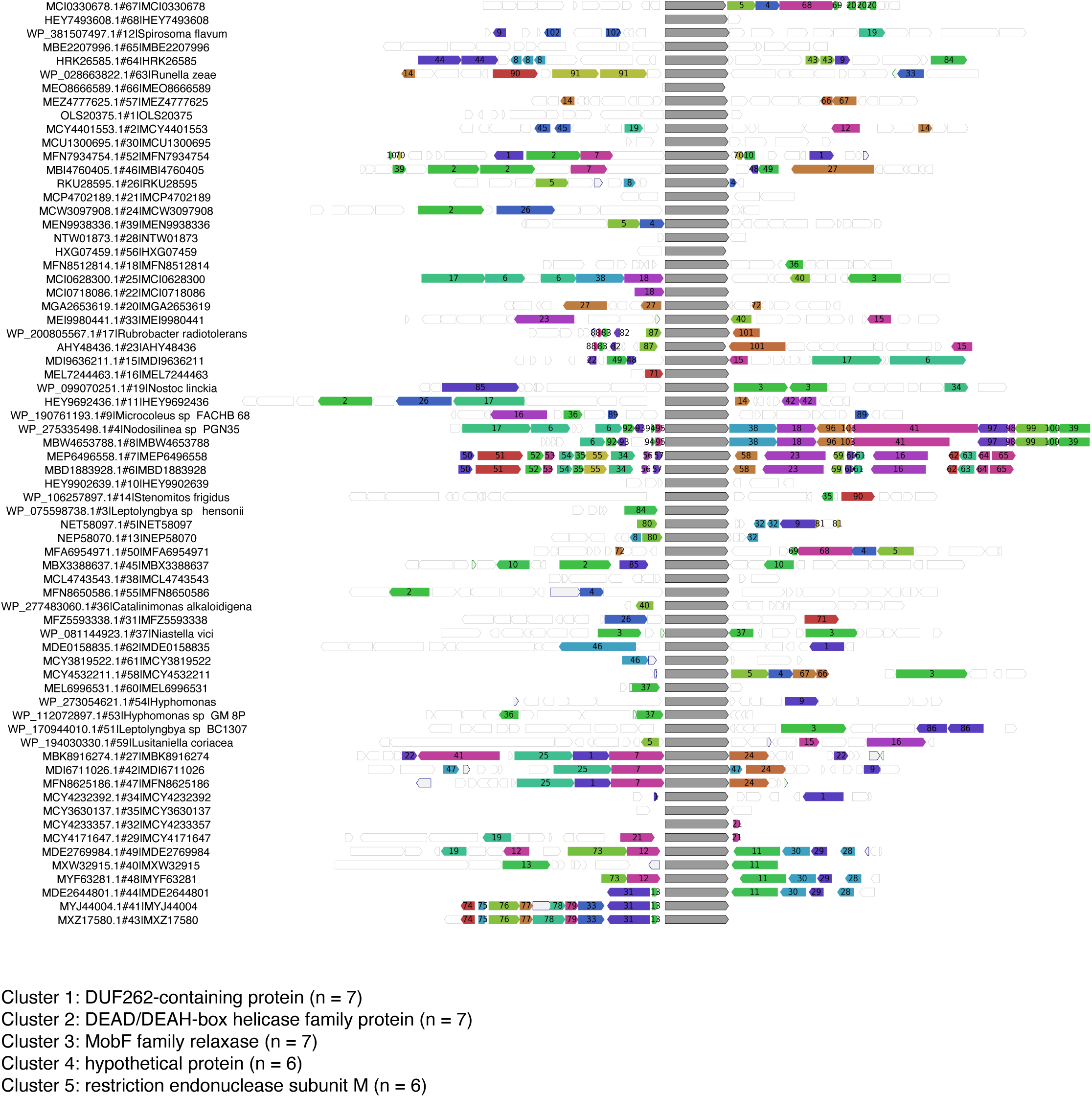
Genomic neighborhood analysis of SLFN-pAgos. FlaGs2 analysis of SLFN-pAgo genomic neighborhoods. SLFN-pAgo genes are shown as grey arrows. For each SLFN-pAgo gene, ten upstream and ten downstream genes were retrieved and clustered according to protein homology. Each cluster was assigned a number inversely proportional to its conservation score; lower numbers therefore indicate proteins more frequently found adjacent to SLFN-pAgos. RzAgo corresponds to RefSeq accession WP_028663822.1. Descriptions of the five most frequently observed flanking gene clusters are shown.

**Extended Data Figure 3.**
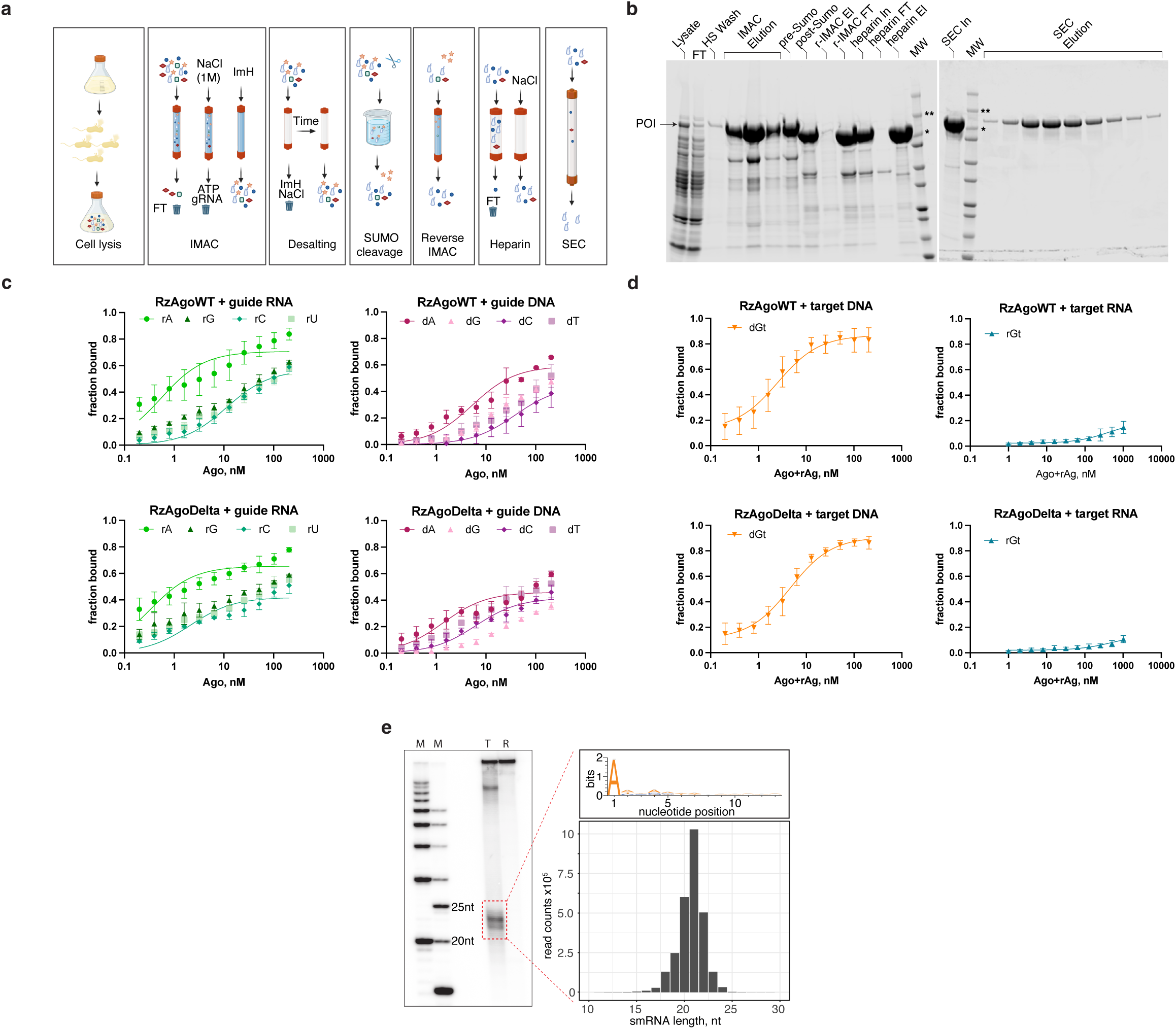
Purification of RzAgo and analysis of its nucleic acid-binding properties in vitro and in vivo. **a,** Schematic of the optimized RzAgo purification workflow (see Methods). **b,** Representative SDS–PAGE analysis showing RzAgo purity and integrity at the purification steps shown in **a**. Protein samples were separated on 4–20% SDS–PAGE gels and visualized with One-Step Blue Protein Gel stain (Biotium). **c,** Binding isotherms from dot-blot assays measuring dissociation constants (*K*d) for RNA or DNA guide binding to wild-type (WT) RzAgo or a Schlatpase-deletion mutant. Guides containing each of the four possible 5′ nucleotides were tested. **d,** Binding isotherms from dot-blot assays measuring *K*d values for RNA or DNA target binding to WT RzAgo or the Schlatpase-deletion mutant. Data in **c** and **d** are mean ± s.d. from *n* = 3–4 independent experiments. **e,** Left, representative denaturing PAGE image of radiolabeled nucleic acids associated with RzAgo in its native host, *R. zeae*. M, ssDNA marker; T, total nucleic acid fraction; R, fraction after RNase A treatment. Right, nucleotide logo and read-length distribution of small RNA guides excised from the gel shown on the left.

**Extended Data Figure 4.**
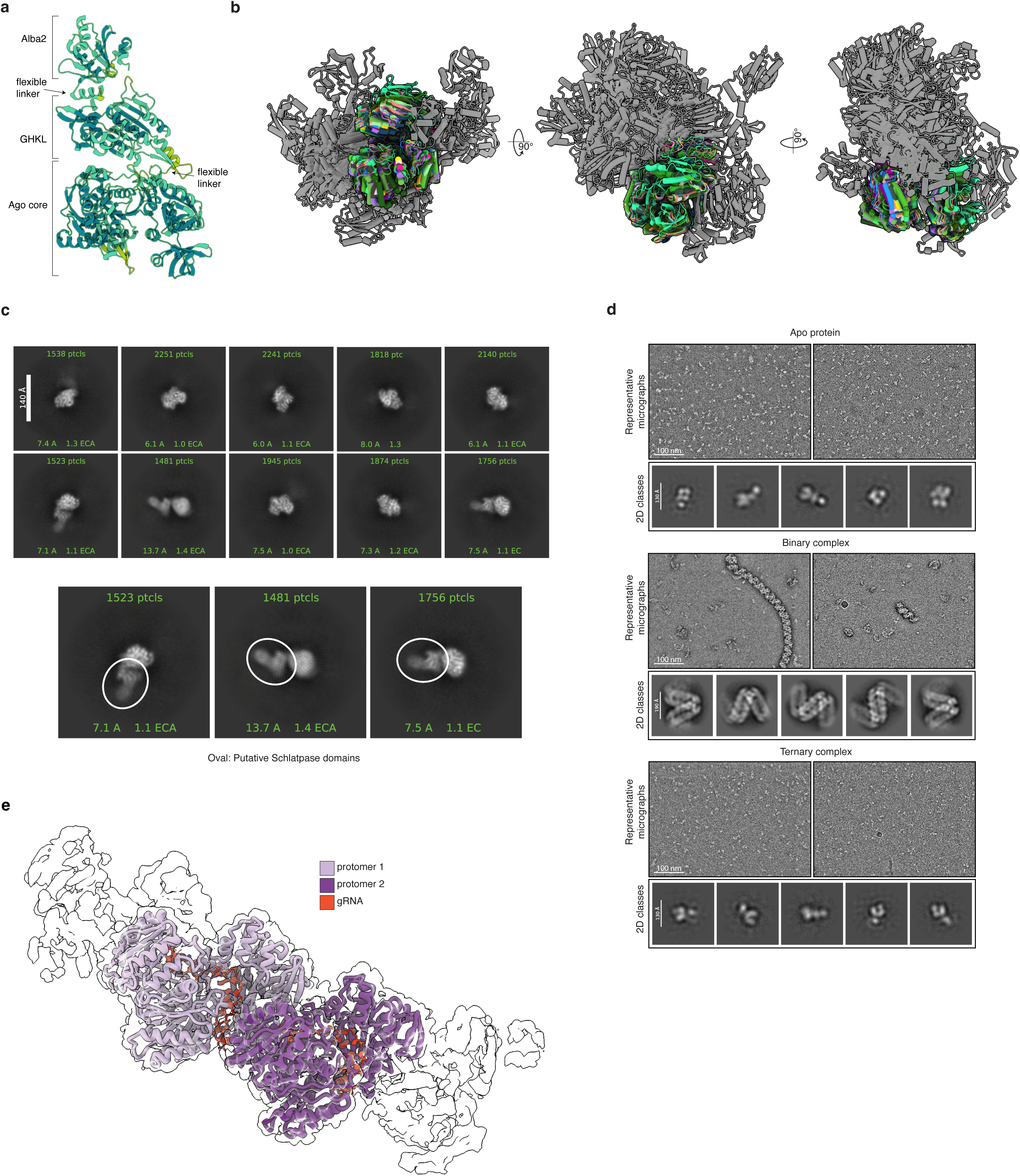
Structural and EM evidence for conformational variability of the Schlatpase supradomain. **a,** AlphaFold3 model of monomeric apo RzAgo. Major domains are indicated, including two linker regions connecting the SLFN_alba2 domain to the GHKL ATPase domain and the GHKL ATPase domain to the pAgo core. **b,** Structural alignment of 19 RzAgo models generated using AlphaFold2 with subsampled multiple sequence alignments to sample alternative conformations. Structures were aligned on the pAgo core domains, which are colored, whereas the Schlatpase supradomain, shown in grey, adopts different orientations relative to the pAgo core. **c,** Top, selected representative 2D class averages from the apo RzAgo cryo-EM dataset. Bottom, enlarged views of three classes highlighting diffuse density, outlined by ovals, likely corresponding to the Schlatpase supradomain. **d,** Negative-stain EM analysis of apo RzAgo, guide RNA-bound RzAgo binary complex and guide RNA–target DNA-bound RzAgo ternary complex. For each condition, two representative raw micrographs and selected representative 2D class averages are shown. Scale bars are indicated. **e,** Cryo-EM structure of the RzAgo filament obtained without crosslinking, showing two consecutive protomers within a low-contour density map shown as a transparent surface.

**Extended Data Figure 5.**
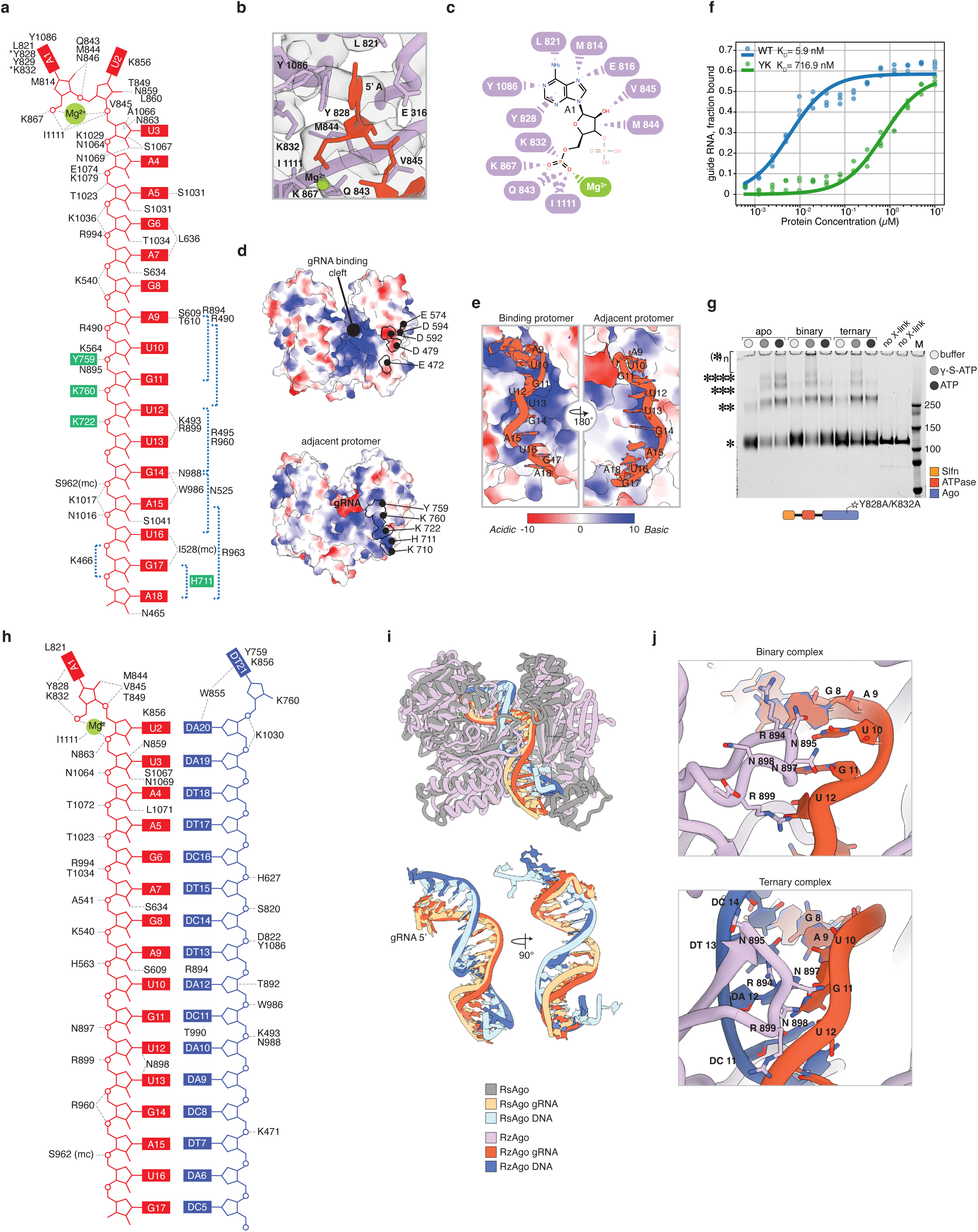
Additional structural and biochemical characterization of the guide RNA pathway in RzAgo binary and ternary complexes. **a,** Contact map showing interactions between RzAgo residues and the guide RNA. Residues from the adjacent RzAgo protomer involved in guide binding are highlighted with green boxes. Blue dotted brackets indicate regions of the guide RNA for which residue assignment is uncertain owing to limited local cryo-EM resolution. **b,** Close-up view of protein–RNA contacts within the 5′-nucleotide-binding pocket of the RzAgo MID domain. **c,** Schematic of interactions between RzAgo residues and the adenine at position 1 of the guide RNA. **d,** Top, electrostatic surface potential of protomer N within the pAgo core filament, revealing a negatively charged surface patch at the position otherwise occupied by the N domain. Bottom, electrostatic surface potential of the adjacent protomer, N−1, revealing a positively charged surface patch that contacts the guide RNA and contributes to the guide-binding tunnel of protomer N. **e,** Electrostatic surface potential of the pAgo core-domain interface within the filament, centred on the 3′ end of the guide RNA and shown from the perspective of the binding protomer (left) and the adjacent protomer (right). **f,** Guide RNA-binding isotherms measured by DRaCALA for WT RzAgo and the Y828A/K832A double mutant (YK). Points show data from three independent experiments, and lines indicate fits to the Langmuir binding model. Dissociation constants (*K*d) are indicated. **g,** Crosslinking SDS–PAGE analysis of oligomerization of the RzAgo YK mutant in apo, binary and ternary states. Positions of monomers, dimers, trimers and tetramers are indicated. no X-link, non-crosslinked control; M, molecular weight marker; γ-S-ATP, non-hydrolysable ATP analogue. **h,** Contact map showing interactions between RzAgo residues, guide RNA and fully complementary target DNA. Guide RNA is shown in red. **i,** Top, structural alignment of the RsAgo ternary complex (PDB 6D8P, grey) with the RzAgo ternary complex (pink). Bottom, two orthogonal views of aligned guide RNA–target DNA heteroduplexes from the RzAgo and RsAgo ternary complexes, showing highly similar duplex conformations. **j,** Structural comparison of the orientation of a PIWI-domain loop, residues 891–903, in the RzAgo binary complex (top) and ternary complex (bottom).

**Extended Data Figure 6.**
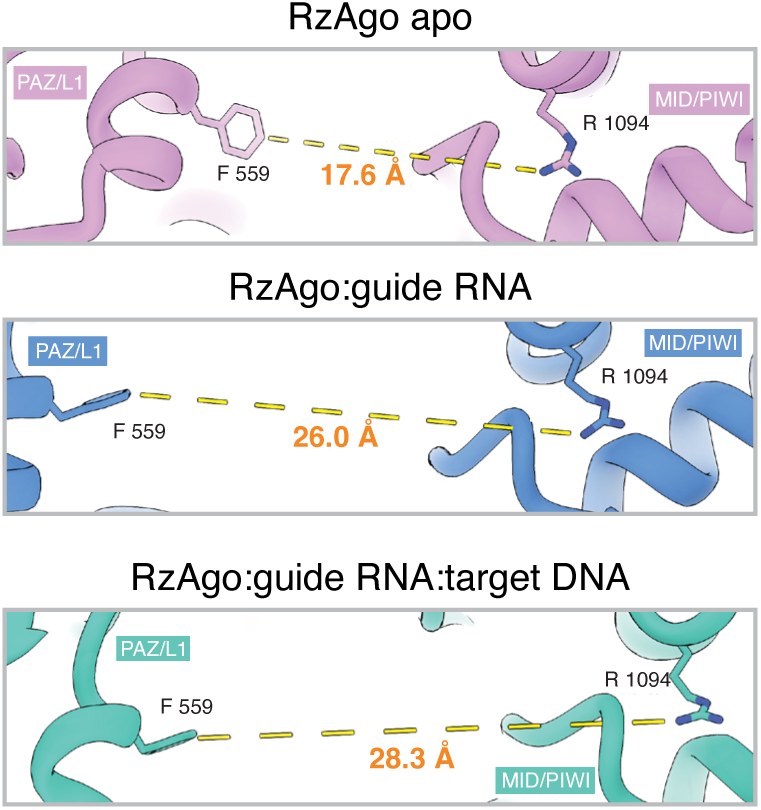
Width of RNA guide binding cleft in apo, binary and ternary RzAgo complexes. Distance between the PAZ/L1 and MID/PIWI lobes of RzAgo in the apo, binary and ternary states, measured between residues F559 in the PAZ/L1 lobe and R1094 in the MID/PIWI lobe. Protein elements from the apo, binary and ternary structures are shown in pink, blue and green, respectively. Key amino acid residues are labelled.

**Extended Data Figure 7.**
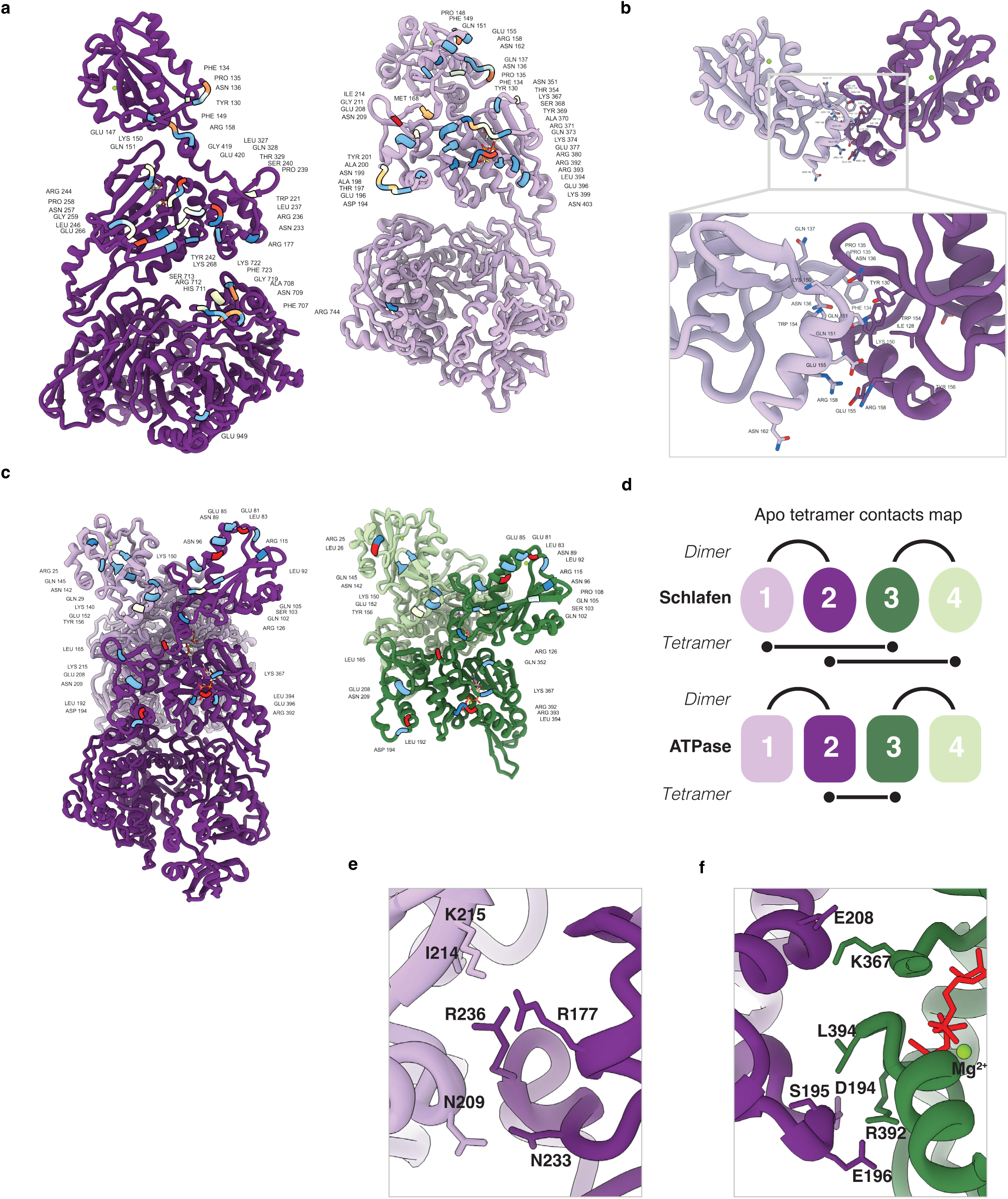
Additional structural characterization of the RzAgo tetramer. **a,** RzAgo monomers adopt two distinct conformations within a dimer: protomer A, shown in light purple, and protomer B, shown in dark purple. Amino acid residues involved in formation of the intradimer interface are colored and labelled. **b,** Detailed view of contacts between two interacting helices at the C-terminal base of the RzAgo SLFN domains within a dimer. **c,** Two RzAgo dimers related by pseudo-C2 symmetry assemble into the tetrameric complex: dimer pair A–B, shown in light and dark purple, and dimer pair A′–B′, shown in light and dark green. The coloring scheme is the same as in Fig. 4f. Amino acid residues involved in formation of the interdimer interface are colored and labelled. **d,** Simplified schematic summarizing domain–domain contacts between RzAgo protomers within the tetramer. **e,** Intersubunit contacts between two RzAgo monomers within a dimer mediated by the N-terminal subdomain. **f,** Interdimer contacts formed by the N-terminal subdomain.

**Extended Data Figure 8.**
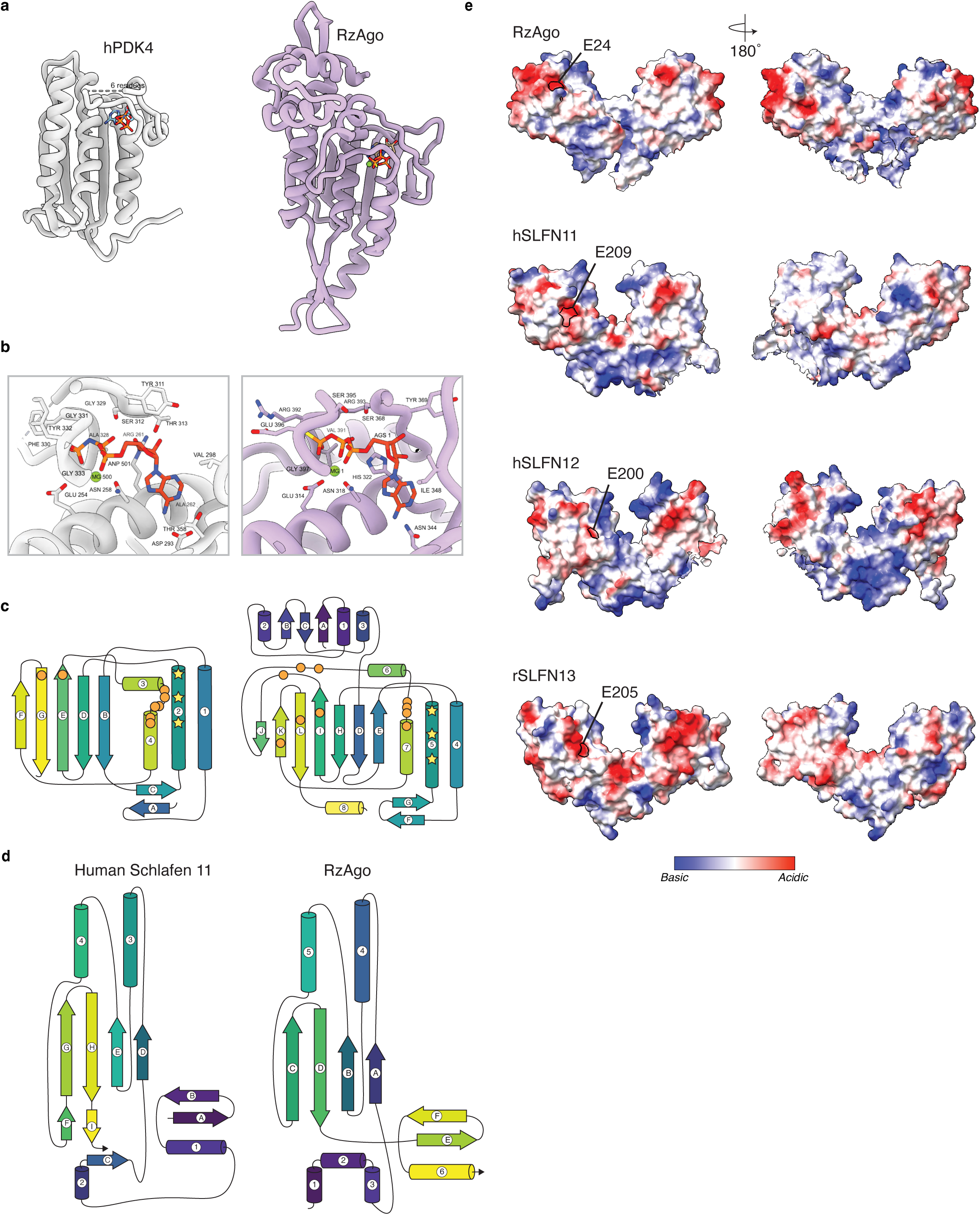
Structural comparison of RzAgo GHKL ATPase and SLFN_alba2 domains with eukaryotic counterparts. **a,** Structural comparison of the GHKL ATPase domain from human PDK4 (PDB 2E0A; left, grey) and RzAgo (right, light purple). **b,** Close-up views of the ATP-binding sites in human PDK4 (left) and RzAgo (right). **c,** Secondary-structure diagrams of the GHKL ATPase domains of human PDK4 (left) and RzAgo (right). **d,** Secondary-structure diagrams of the C-lobe of the SLFN domain from human SLFN11 (left) and the SLFN_alba2 domain of RzAgo (right). In mammalian SLFN proteins, the complete SLFN module consists of an N-lobe, linker and C-lobe; only the C-lobe shares homology with the SLFN_alba2 domain present in RzAgo (see Fig. 5d). **e,** Electrostatic surface potential maps comparing the arrangement of SLFN_alba2 domains in the RzAgo apo tetramer (top row; one dimer is shown) with full-length SLFN domains from mammalian SLFN proteins: human SLFN11, human SLFN12 and rat SLFN13. Two orthogonal views are shown. **f,** Space-filled model of the asymmetric three-protomer SLFN-domain cluster in the RzAgo filament, colored by protomer. The green dashed outline marks the position of the SLFN dimer observed in the apo tetramer. In this arrangement, the third SLFN domain sterically occludes the putative substrate-binding cleft.

**Extended Data Figure 9.**
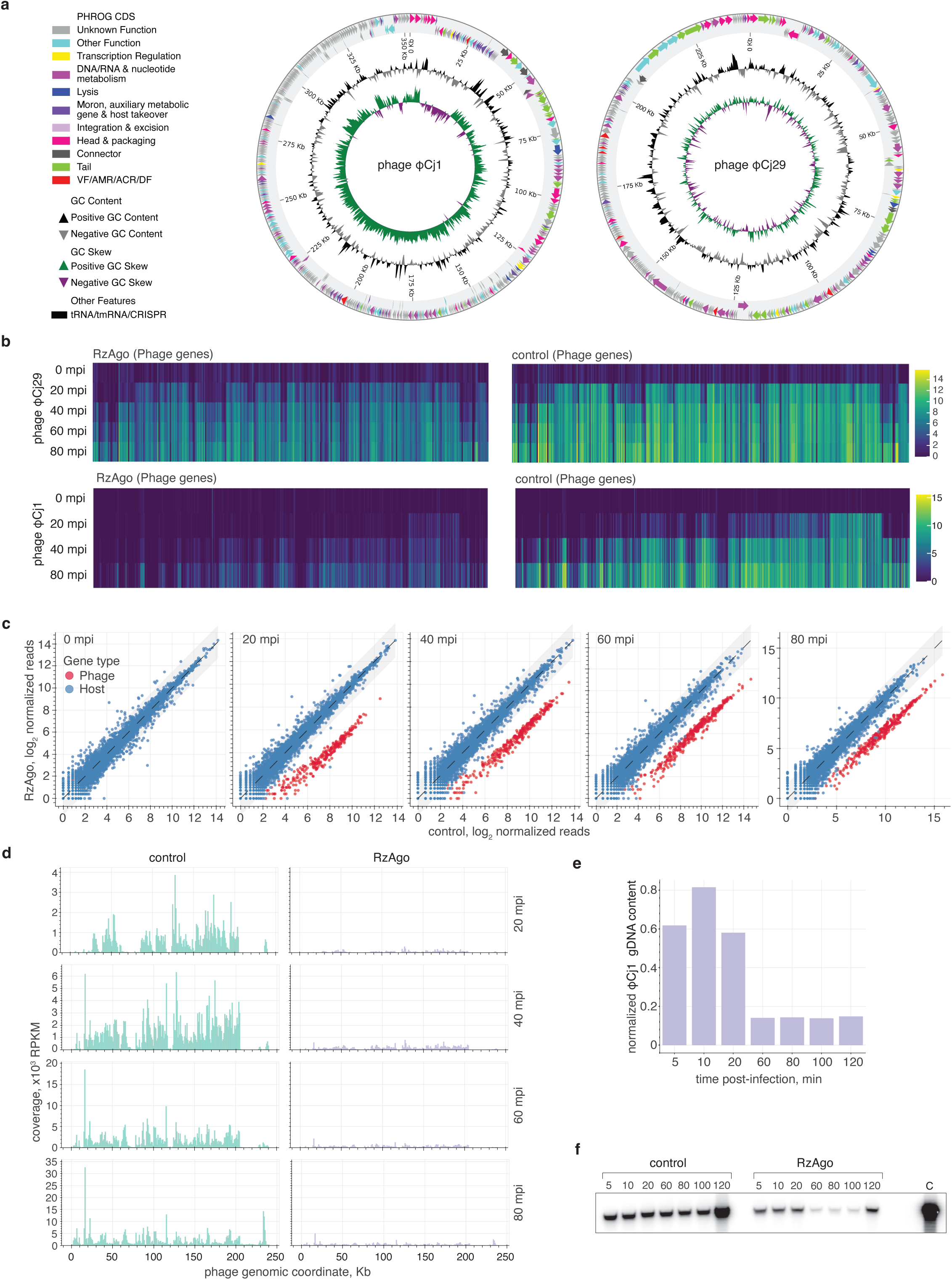
Additional data illustrating the effect of RzAgo on phage genome replication and transcription. **a,** Circular genome maps of two jumbo phages, φCj1 and φCj29, infecting *F. johnsoniae* and assembled in this study. Genes are colored according to the eleven PHROG groups. Tracks showing GC content and GC skew are indicated. **b,** Heat maps showing log₂-transformed transcript abundance for φCj29 (top row) and φCj1 (bottom row), comparing cells expressing RzAgo (left column) with control cells carrying an empty vector (right column). **c,** Scatter plot of total RNA-seq data, shown as log₂-transformed TMM-normalized reads, illustrating gene-expression changes during φCj29 infection. Blue dots represent host genes, and red dots represent phage genes. The shaded grey region indicates a ±1.5 log₂ fold-change interval. **d,** Coverage plots showing the distribution of total RNA-seq reads, normalized as RPKM, across the φCj29 genome at 20, 40, 60 and 80 min post-infection. **e,** Bar plot showing qPCR analysis of φCj1 genomic DNA levels in cells expressing RzAgo at different time points post-infection, normalized to cells carrying an empty-vector control. **f,** Southern blot of total genomic DNA isolated from cells expressing RzAgo or carrying an empty vector at different time points after infection with phage φCj1. The blot was probed using a fragment corresponding to an approximately 3-kb region of the φCj1 genome; mpi – minutes post-infection.

**Extended Data Figure 10.**
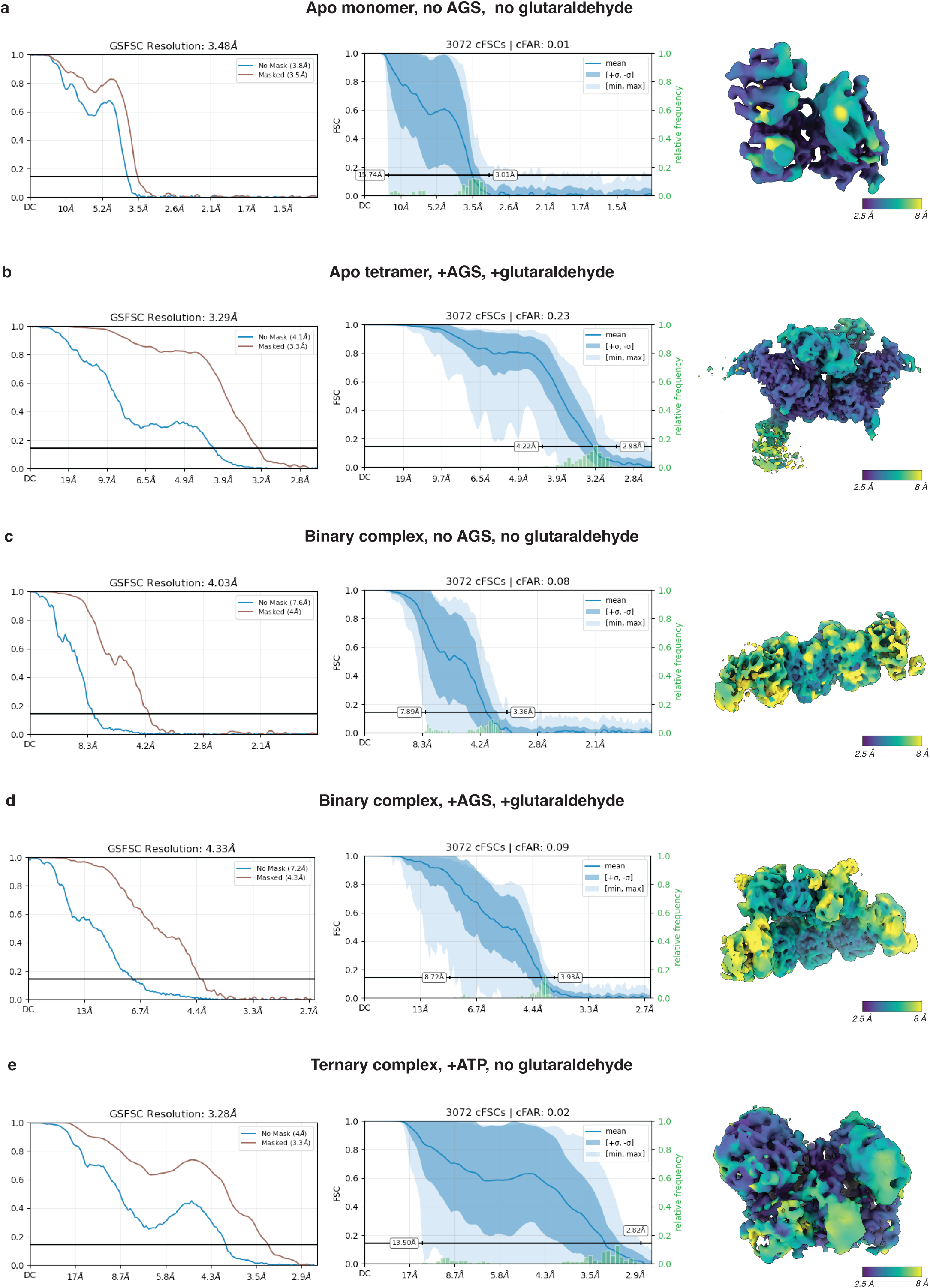
Cryo-EM map quality assessment. Cryo-EM map validation metrics for five reconstructed RzAgo structures. For each reconstruction, gold-standard Fourier shell correlation (GSFSC) plots and directional cFAR plots are shown together with the corresponding cryo-EM density map colored by local resolution and displayed using the Viridis color map (purple, 2.5 Å; yellow, 8 Å). **a,** Apo RzAgo monomer reconstructed in the absence of γ-S-ATP and glutaraldehyde. **b,** Apo RzAgo tetramer reconstructed in the presence of γ-S-ATP and mild glutaraldehyde crosslinking. **c,** Binary RzAgo–guide RNA complex reconstructed in the absence of γ-S-ATP and glutaraldehyde. **d,** Binary RzAgo–guide RNA complex reconstructed in the presence of γ-S-ATP and mild glutaraldehyde crosslinking. **e,** Ternary RzAgo–guide RNA–target DNA complex reconstructed in the presence of ATP and in the absence of glutaraldehyde.

## Notes

### Competing Interest Statement

The authors have declared no competing interest.

